# Molecular plasticity of the native mouse skeletal sarcomere revealed by cryo-ET

**DOI:** 10.1101/2020.09.13.295386

**Authors:** Zhexin Wang, Michael Grange, Thorsten Wagner, Ay Lin Kho, Mathias Gautel, Stefan Raunser

**Affiliations:** Department of Structural Biochemistry, Max Planck Institute of Molecular Physiology, Otto-Hahn-Strasse 11, 44227, Dortmund, Germany; The Randall Centre for Cell and Molecular Biophysics, School of Basic and Medical Biosciences, Kings College London BHF Excellence Centre, New Hunt’s House, Guy’s Campus, London SE1 1UL, UK

**Author notes:** The authors contributed equally (Zhexin Wang, Michael Grange).

## Abstract

Sarcomeres are the force-generating and load-bearing devices of muscles. A precise molecular understanding of how the entire sarcomere is built is required to understand its role in health, disease and ageing. Here, we determine the *in situ* molecular architecture of vertebrate skeletal sarcomeres through electron cryo-tomography of cryo-focused ion beam-milled native myofibrils. The reconstructions reveal the three-dimensional organisation and interaction of actin and myosin filaments in the A-band, I-band and Z-disc and demonstrate how α -actinin cross-links antiparallel actin filaments to form a mesh-like structure in the Z-disc at an unprecedented level of molecular detail. A prominent feature is a so-far undescribed doublet of α-actinin cross-links with ∼ 6 nm spacing. Sub-volume averaging shows the interaction between myosin, tropomyosin and actin in molecular detail at ∼ 10 Å resolution and reveals two coexisting conformations of actin-bound heads. The flexible orientation of the lever arm and the essential and regulatory light chains allow the two heads of the “double-headed” myosin not only to interact with the same actin filament but also to split between two actin filaments. Our results provide new insights into the conformational plasticity and fundamental organisation of vertebrate skeletal muscle and serve as a strong foundation for future *in situ* investigations of muscle diseases.

## Introduction

Skeletal muscle is an essential tissue required for efficient movement in vertebrates. In humans, dysfunctional muscle function leads to a broad variety of physiological disorders ranging from common muscle cramps to severe myopathies^1,2^. The cellular unit of muscles is a muscle fibre. Individual muscle fibres can be very large syncytia of up to several centimetres in length and are comprised of the force-generating and load-bearing devices of muscles called sarcomeres. Two Z-discs border the sarcomere at its ends, anchoring the barbed (+) ends of actin (thin) filaments. Their pointed (-) ends face towards centrally positioned bipolar myosin (thick) filaments. The myosin filaments are stably linked to actin filaments by titin^3^. The basis of the mechanism of skeletal muscle action is the cyclical interaction of actin and myosin, which leads to relative movement between thin and thick filaments and is the primary driver of force generation within the sarcomere^4,5^.

The current understanding of actin-myosin interactions and the function and mechanism of other essential sarcomeric proteins is derived from atomic structures of *in vitro* reconstituted components^6–10^, necessarily lacking important information in their biological context, or ensemble information derived from X-ray diffraction of muscle fibres^11^. Reconstructions of insect sarcomeres using conventional electron tomography^12–14^ and freeze-substituted samples^15^ gave insights into the three-dimensional structure of sarcomeres. However, the studies were limited to low resolution and did not allow an understanding of the molecular detail underpinning their organisation. Direct structural analyses on the native organisation of vertebrate sarcomeres are limited to two-dimensional projection images^16–18^ or three-dimensional reconstructions of chemically stained and treated myofibrils with unresolved cross-bridges^19^. Here, using electron cryo-tomography (cryo-ET), we present the three-dimensional structure of a mouse psoas sarcomere in the rigor state with a comprehensive description of the organization and interaction of sarcomeric proteins at an unprecedented level of molecular detail.

### Three-dimensional organisation of the sarcomere

The major obstacles in obtaining molecular structural details of a sarcomere using conventional methods lie in artefacts arising from fixation and staining methods, the diamond blade during sectioning or from the shrinkage of plastic sections during imaging^20^, impeding the preservation of high-resolution features. To circumvent these artefacts, we vitrified mouse psoas myofibrils in the rigor state (i.e. without ATP) by plunge-freezing in liquid ethane, preserving the hydrogen bond network and fine ultrastructure, and subsequently prepared thin lamellae using cryo-focus ion beam milling (cryo-FIB)^21^ (Methods). The lamellae were 30 – 150 nm thin and ideally suited for electron cryo-tomography (cryo-ET) (Figure S1).

The vitrified myofibrils show an intact sarcomere ultrastructure, including apparent Z-discs (actin filaments and their crosslinks), I-bands (only actin filaments), A-bands (zone of overlapping actin and myosin filaments), M-bands (myosin filaments and their crosslinks), as well as organelles such as mitochondria and sarcoplasmic reticulum localised between laterally adjacent sarcomeres (Figure 1a, S1). Importantly, the tomograms include structures that could previously not be visualized *in situ*, in particular myosin double heads, forming regular cross-bridges that resemble an “arrow-head”, thin myosin tails that are mainly composed of coiled-coil structures^22^ protruding from the thick filament, details of the M-band, Z-disc and troponin complexes (Figure 1b-d).

**Figure 1.**
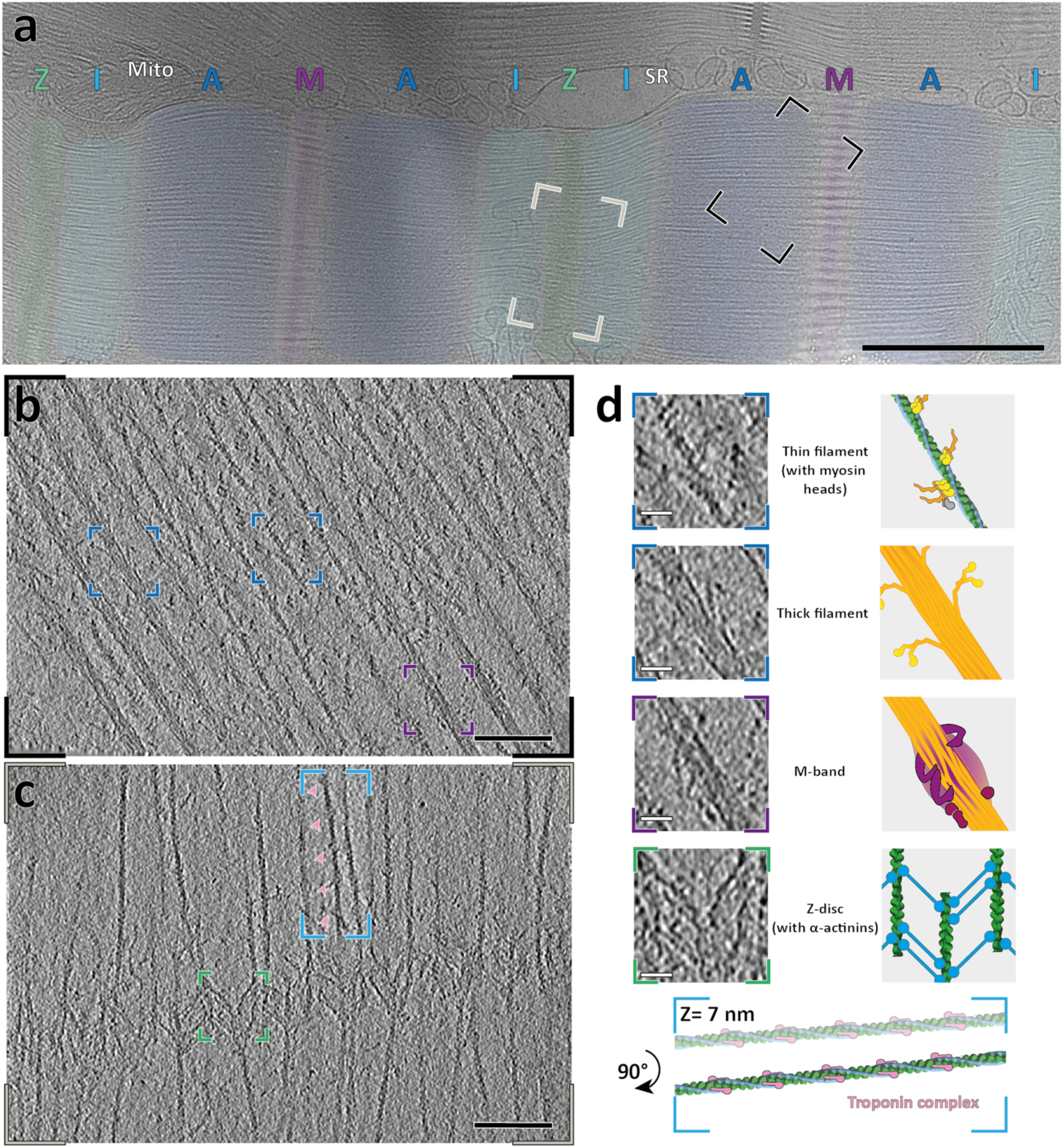
Isolated mouse skeletal myofibrils imaged using electron cryo-tomography. **a**, Projection image of mouse skeletal muscle exhibits all the hallmarks of a sarcomere, with Z-disc, I-, M-, and A-bands clearly visible (green, light blue, dark blue and purple, respectively). **b**, Slice through an electron cryo-tomogram acquired on a region with stretching from an A-to an M-band (black inset in (**a**) shows a representative position). In this region, it is possible to discern myosin heads bound to the thin filament, the branching of myosin tails from the thick filament within the A-band (dark blue insets) and a widening of the thick filaments at the M-band due to the presence of obscure protein densities (purple inset). **c**, Slice through an electron cryo-tomogram acquired on a region stretching from a Z-disc to an I-band (grey inset in (**a**) shows a representative position). The arrow-tail-feather-like arrangement of α-actinin molecules cross-bridging thin filaments can be clearly observed in a zig-zag manner (green inset), while the thin filaments in the I-band have regularly-spaced nodes that correspond to the troponin-complex (light pink arrow heads, blue inset). **d**, Larger view of insets described in (**a-c**), with cartoon depictions of densities. Scale bars, 1 µm (**a**), 100 nm (**b, c**) and 20 nm (**d**).

It is well-known from cross-sectional views of myofibrils that thin and thick filaments are hexagonally arranged in the A-band^16^. In orthogonal views of our tomograms, we can clearly discern this hexagonal pattern in the A-band, demonstrating that our preparations are consistent with previous studies (Figure S2a-c). One thin filament is surrounded by three thick filaments and three other thin filaments at a distance of around 26 nm (Figure S2). Thick filaments form triangular patterns with an averaged inter-thick-filament distance of 45 nm (Figure S2a-c). This arrangement of thick filaments is maintained from the A-band to the M-band where thin filaments are missing (Figure S2d-f).

### In situ actin-myosin interactions in a vertebrate sarcomere

Segmentation of the tomograms enabled us to unequivocally characterise the 3D arrangement of thin and thick filaments and the cross-bridges within a sarcomere (Movie S1). To increase the level of observable detail that we could resolve with our data, we employed sub-volume averaging to determine the structure of the thin and thick filament and cross-bridges. The reconstruction of the thick filament proved to be difficult due to its heterogeneity, flexibility and an absence of obvious symmetry. Nevertheless, we obtained a low-resolution reconstruction showing the principle architecture of the core of thick filaments that we could use for the following analysis (Figure S3).

The three-dimensional reconstruction of the thin filament and cross-bridges could be resolved to a higher resolution. Segmentation of the tomograms and 3D classification of all sub-volumes of the cross-bridges indicated that the two heads from the same myosin molecule, known as double-heads, mostly bind to two neighbouring actin subunits (Figure S4d). Hence, we merged multiple classes after modifying alignment parameters to obtain - to the best of our knowledge - the first structure of a native thin filament with myosin double-heads bound (Methods, Figure 2a). The reconstruction reached a resolution of 10.2 Å and we could clearly assign tropomyosin and the domains of actin and of the myosin heads and necks, including the essential chains (ELC) and part of the regulatory light chains (RLC) (Methods, Figure 2a, Figure S4).

**Figure 2.**
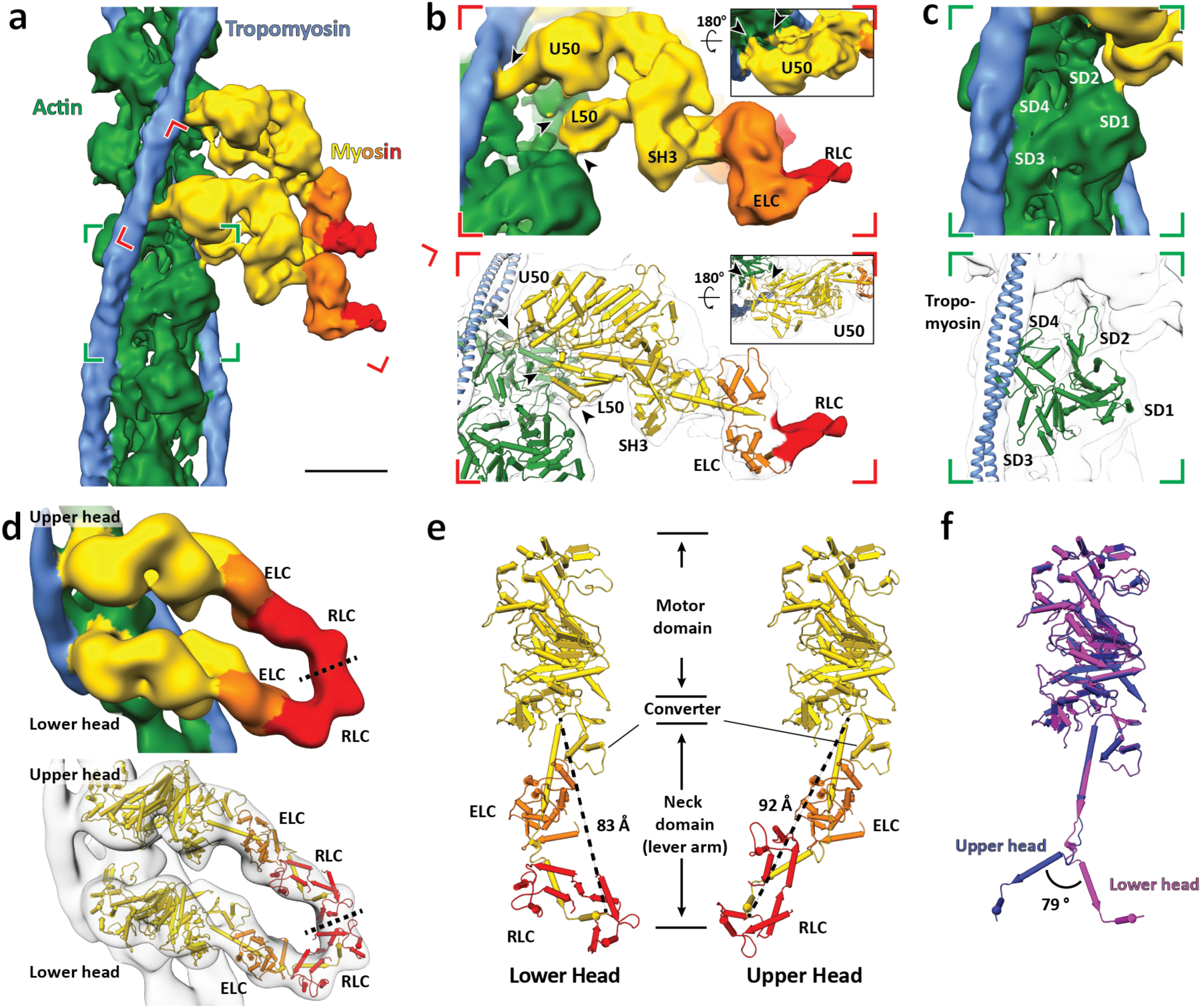
Sub-tomogram averaging of thin filaments reveals the interaction of a double-head myosin with the thin filament and two conformations of light chain domains within a double-head. **a**, Surface view of the *in situ* actomyosin structure at 10.2 Å, showing the actin filament (green), tropomyosin (blue) and a pair of myosin head and neck domains (yellow) with the essential light chains (orange) and regulatory light chains (red). Scale bar, 5 nm. **b**, Close-up view of the lower myosin head and homology models based on PDBs: 3I5G, 5JLH and 6KN8 fitted into the density map. Domains such as the upper 50 kDa (U50), lower 50 kDa (L50), SH3, and essential light chains (ELC) can be allocated in the map. Only part of the regulatory domain (RLC) was resolved in this structure. Arrow heads indicate the interaction interfaces between actin and myosin at loop 4, helix-loop-helix motif, loop 3 of myosin from top to bottom. Arrow heads in the inset depict the interaction interfaces at the cardiomyopathy loop and loop 2 from left to right. **c**, Close-up view of an actin subunit and the structural model fitted into the EM map showing the four subdomains of an actin subunit (SD1-4). **d**, Surface view of the structure of a complete myosin double-head including regulatory light chains determined from averaging re-extracted shifted sub-volumes (see Figure S4). The two RLCs (homology model based on PDB: 3I5G) were rigid-body fitted into the map. Their interface is indicated by a dotted line. **e**, Comparison between the lower and upper heads within one myosin double head, which shows two different conformations in the lever arm that interacts with RLC and ELC. Lengths of the lever arms were measured between G772 and L844. **f**, Alignment of the lower (purple) and upper (blue) heads heavy chain, showing two different kinks between the ELC-binding region and the RLC-binding region.

We calculated a homology model using the rigor cytoplasmic actomyosin cryo-EM structure (PDB: 5JLH)^7^ and the crystal structure of the S1 fragment from squid muscle (PDB: 3I5G)^23^ as template and fitted it into the density map using rigid body fitting (Figure 2b-c). We chose this myosin structure, because there are no crystal structures of a complete vertebrate myosin S1 fragment in the rigor state available. The model fits well into the density, demonstrating that the *in vitro* structures resemble well the structures determined *in situ*. At this resolution, both myosin heads bind to the actin filament in structurally identical manner with close interactions formed between actin subunits and myosin as well as tropomyosin (Figure 2b, c). The ELCs bind to the upper part of the lever arm (residues L788-L800) as in the crystal structure. The density threshold corresponding to the RLCs and RLC binding domain after averaging is considerably lower than the rest of the structure, indicating a higher flexibility or variability in this region.

To improve the quality of the reconstruction in the region of the RLC, we shifted the centre of sub-volume averaging from the thin filament to the myosin head. Through further classification, we determined a structure of the complete double-head myosin S1 fragment, including the two heavy chains that are bound to actin, two ELCs and two RLCs (Figure 2d). The S2 fragment that tethers myosin heads to the thick filament was not resolved in this structure, likely due to the different positions and angles it takes in the sarcomere.

Although a homology model of the myosin motor domain and ELC of the squid myosin crystal structure (PDB: 3I5G) fits well into our density, the RLCs and the lower part of the lever arm (residues R811-L844) had to be fitted separately (Figure 2d). Compared with its conformation in the crystal structure, the lever arm is strongly kinked between the ELC and RLC binding regions (residues M801-E810), resulting in a bent conformation (Figure 2e). Interestingly, the kink in the upper head is exactly in the opposite direction compared to the one in the lower head, bringing the RLCs of the two heads in close proximity (Figure 2f). This is also consistent with previous cross-linking and FRET experiments, which showed a small distance between the two RLCs in a double head^24–26^.

This arrangement, which is probably stabilized by interactions between the ELCs and RLCs, allows for the simultaneous binding of the two heads to actin and at the same time brings their necks close enough together to continue with the coiled-coil structure of the joint S2 fragment. In addition, both lever arms have a similar length. This is important, because the length of the lever arm determines the step size of myosin heads on actin^27^.

In this context, it is interesting that, the hinge in the lever arm together with a hook (residues W824-W826 in molluscan myosin) have previously been shown to be important for regulated myosins^28,29^. In this case, the light chains control myosin activity even in the absence of actin^30^ and the presence of Ca^2+^ determines the stability of the ELC/RLC interface. Different ELC/RLC interfaces were also suggested from models of myosin in the relaxed state, where the heads fold back to the thick filament^31^. Notably, the arrangement of the ELC/RLC of the lower head in our model resembles that in the “blocked” head of the off-state myosin while the upper head ELC/RLC is similar to the “free” head (Figure S5). This suggests that there are only two possible angles for the kink in the lever arm, which are likely to be determined by the interaction of the ELCs and RLCs. This interaction probably stabilizes the two conformations to rigidify the lever arm, which is needed for the proper transmission of the force of the power stroke^32^.

### Cross-bridges in the A-band

In order to identify the proportion of myosin heads that bind to thin filaments in a rigor state sarcomere, we annotated and isolated the densities corresponding to the cross-bridges as well as thick and thin filaments in the A-band (Figure 3a, S6). Combining the annotated filaments with the structures determined via sub-volume averaging allowed us to produce a molecular map of the thin and thick filaments and attached myosin heads (Figure 3b, S7).

**Figure 3.**
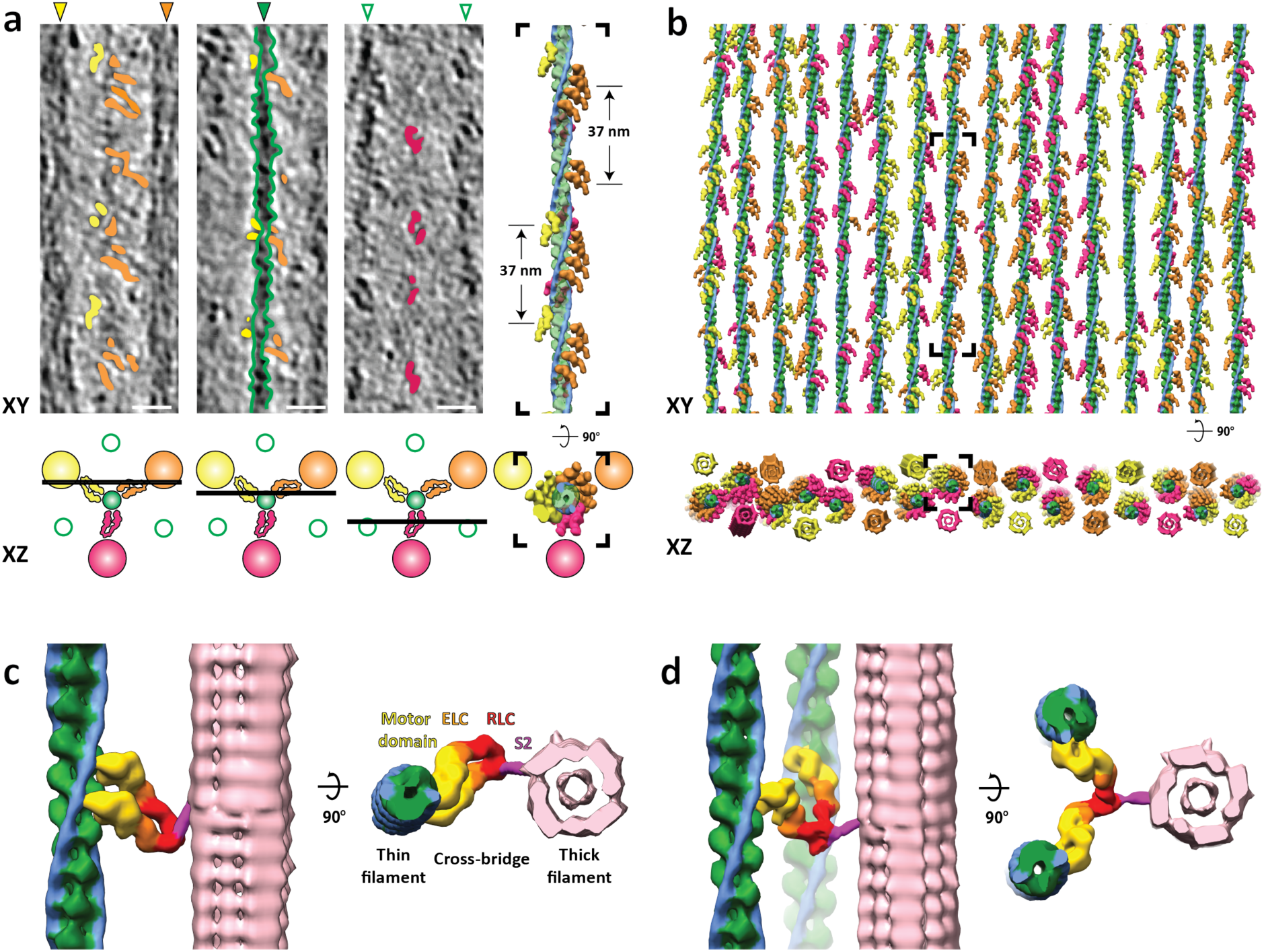
Organisation of the A-band in natively isolated myofibrils shows that myosin heads can adopt two interactions with thin filaments. **a**, XY-slices through the tomogram at three different Z positions, depicted by the cartoons below, which demonstrate the myosin head densities (orange, yellow and magenta) and the thin filament (outlined with green dotted lines) apparent within the tomographic volume. Annotation of these densities results in a volume in which the filtered 3D reconstruction of actomyosin can be fitted, which is shown on the right. The three myosin colours represent the contribution of the myosin head from a respective neighbouring thick filament. Scale bars, 20 nm. **b**, Fitted model of all thin filaments and myosin heads in a tomogram, with the black box depicting the filament shown in (**a**). In the XZ view, there are also the reconstructions of corresponding thick filaments shown. **c**, A typical cross-bridge with a myosin double-head. The model was obtained from fitting maps from sub-volume averaging into annotated densities of the tomogram. The S2 fragment was derived from the crystal structure of the human S2 fragment (PDB: 2FXM), filtered and placed manually according to the annotated densities. **d**, A rare myosin split-head with two heads from the same myosin molecule binding to two different actin filaments. Both heads were fitted with the upper head density shown in Figure 2d. See also Figure S6.

Of the 30 thin filaments annotated containing 5,664 actin subunits, 2,734 myosin heads were fitted into the cross-bridge densities, corresponding to 82.5% of the total number of myosin heads in this region calculated from the theoretical number of myosin heads per given length of the thick filament (6 heads per 14.3 nm, total heads = 3,315). This suggests that not all myosin heads are attached to thin filaments, even in the rigor state.

Similar to sarcomeres from insect muscle, all thin filaments are in helical register (Figure 3b) and cross-bridges appear at regular target regions every ∼37 nm on a thin filament between each pair of thick and thin filaments (Figure 3a, b)^33^. However, in contrast to the “double-chevron” arrangement of cross-bridges in an insect sarcomere, the cross-bridges in a vertebrate sarcomere show a higher variability regarding the distance between each other and tend to appear in clusters (Figure 3a, b and S7). While the most common composition of a cross-bridge is only one myosin double-head (Figure 3c), two consecutive myosin double-heads also appear at many target regions. Occasionally, there are also single heads bound to the thin filament (8.3% of all myosin heads) with their partners not identified within the tomogram. In addition, a rare “split-head” conformation also occurs when the two heads from one myosin binds to two different adjacent thin filaments (10 pairs observed) (Figure 3d, S6). This conformation has been previously suggested^34^ and vaguely indicated by 2D projection images^17^. Our observation provides direct proof of this conformation in three-dimensions in the rigor state vertebrate sarcomere.

To further investigate the variability in myosin binding, we focused on the influence of actin orientation and the distance between thin and thick filaments (Figure S8). We plotted the orientation of the actin subunits that are bound by myosin heads in terms of their relative angles with respect to a fixed orientation perpendicular to the filament axis. The distribution showed three distinctive groups with a Gaussian distribution corresponding to the three thick filaments from which the myosin heads originate. For a specific thick filament, the orientations of myosin-bound actin subunits are mostly confined within a range of ∼120° (Figure S8a). This indicates that myosin heads tend to bind to actin subunits oriented towards the direction of the thick filament from which it originates. This is also demonstrated by the combined myosin binding profile of all 30 thin filaments (Figure S8b, S9). In addition, the footprint map of myosin binding on a thin filament also exhibits the 37 nm hot spot (∼13 actin subunits) of cross-bridge clusters as a consequence of this preferred binding orientation of actin subunits.

Overall, the analysis of myosin heads in the A-band suggests that myosin binding is a stochastic process regulated by physical limitations such as the orientation of actin subunits. Myosin prefers to bind to regions on thin filaments that orient towards the thick filament, creating ∼37 nm periodic cross-bridge clusters, consistent with previous observations in insect muscle^33^ and single-molecule experiments^35^. The S2 fragment, which forms a convex surface with the S1 fragments^36^, provides enough flexibility for a myosin head to bind to a random actin subunit within the range of allowed distance and orientation (Figure S7c). This flexibility makes allowance for the mismatch of the 37 nm actin repeat in thin filaments and the 14.3 nm myosin repeat in thick filaments^37^ and results in the pseudo-regular arrangement of cross-bridges.

### Thin filaments in the I-band

One advantage of our approach is that we can trace single filaments through the sarcomere in 3D, allowing us to follow thin filaments from the A-band through the I-band into the Z-disc. The ordered hexagonal pattern of filaments in the A-band breaks down in the I-band, where the thick filaments end (Figure 4d). Consistent with previous studies presenting cross-section images of the I-band^38^ and contrary to a previous model stating an equally-distanced arrangement of thin filaments along the A-band to Z-disc transition^39^, thin filaments show a transition from a hexagonal pattern to an irregular pattern caused by the lack of cross-bridges in this region (Figure 4d). The extent of displacement from the hexagonal arrangement varies among filaments.

**Figure 4.**
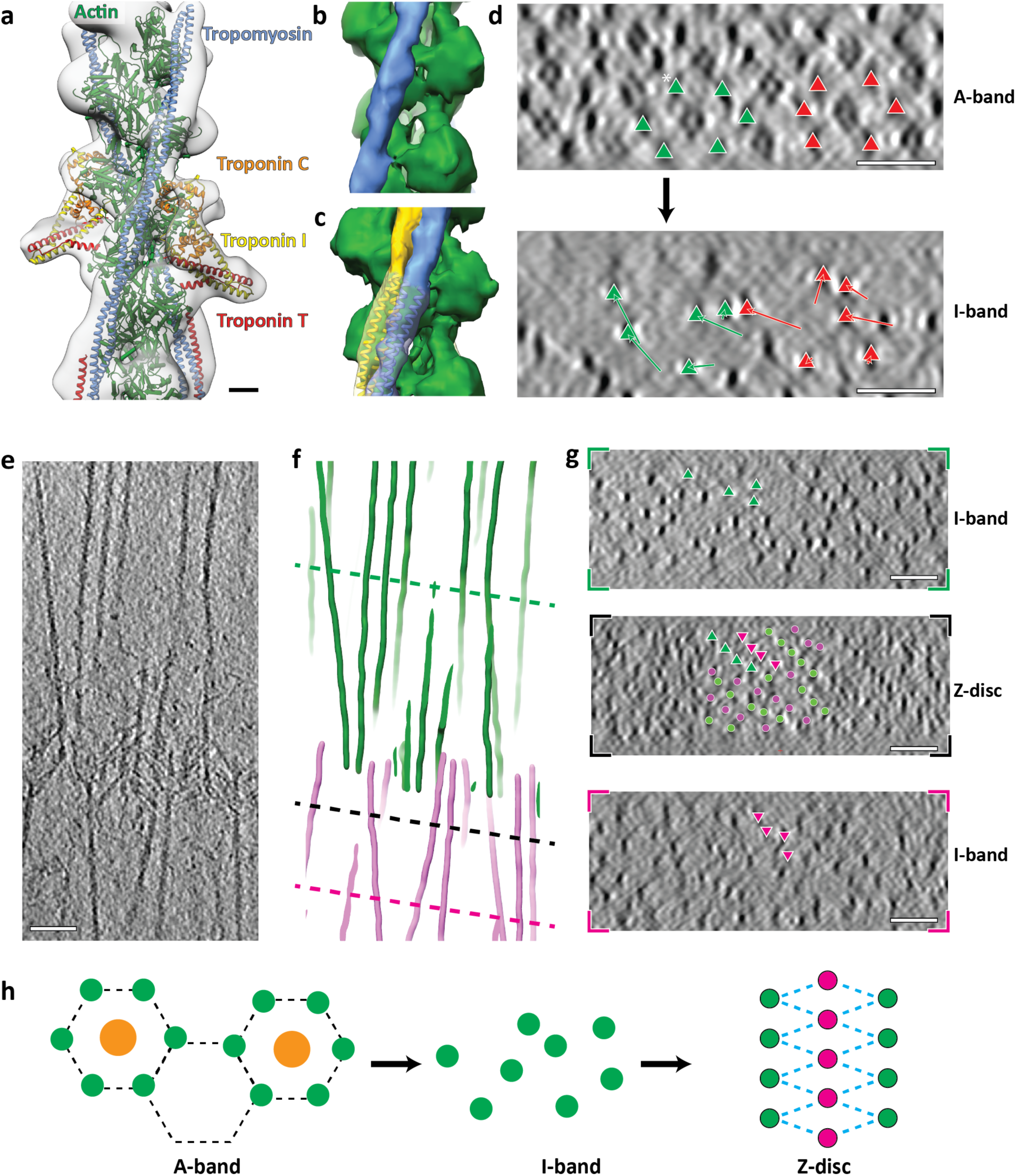
Structure and organisation of thin filaments in the I-band. **a**, Structural model of the thin filament in the I-band fitted into the 3D reconstruction of a complete thin filament including troponin. This homology model is based on the structure of the Ca^2+^-bound cardiac muscle thin filament (PDB: 6KN8). Scale bar, 2 nm. See also Figure S10. **b**, 3D reconstruction of the thin filament in the I-band excluding troponin. The actin filament is depicted in green and tropomyosin in blue. **c**,**d** While tropomyosin takes the C-state in the I-band (blue), it is in the M-state when myosin heads are bound to the thin filament in the A-band (yellow). See also Figure S10. **d**, Cross-section views of a sarcomere highlight the disappearance of the hexagonal pattern of thin filaments during transition from the A-band to the I-band. Two hexagonal units of thin filaments and their corresponding positions in the I-band are indicated by green and orange triangles. The displacement of the filaments from A-band to I-band are shown as arrows. The filament marked with an asterisk moved out of the field of view during the A-I transition. **e**, Slice through a tomogram depicting the Z-disc and two I-bands from two adjacent sarcomeres. **f**, 3D model of thin filaments showing the same region as in (**e**). **g**, Cross-section views of the positions indicated by dotted lines in (**f**), showing the pattern of thin filaments during the I-Z-I transition. Filaments are traced in green triangles from the I-band to Z-disc of the top sarcomere and in magenta triangles from the Z-disc to the I-band of the bottom sarcomere. In the Z-disc image, antiparallel filaments in the centre are labelled with green and magenta dots for better visualisation. **h**, A schematic diagram shows the hexagonal pattern in the A-band, the irregular pattern in the I-band and the rhomboid pattern in the Z-disc. Scale bars, 50 nm (**d-g**).

Because of the absence of myosin in the I-band, troponin complexes are clearly visible with a periodicity of ∼37 nm on the thin filament (Figure 1c, d, S10). We applied sub-volume averaging to determine the structures of the troponin actin-tropomyosin complex and the actin-tropomyosin complex in the I-band at resolutions of 19.8 Å and 10.3 Å (Figure 4a, b, S10), respectively. We then calculated a homology model using the *in vitro* cardiac muscle thin filament cryo-EM structure of the Ca^2+^-bound state as a template (PDB: 6KN8)^40^ and fitted it into the density map using rigid body fitting. The model fits well into the reconstruction of both structures, demonstrating that the thin filament is in the on-state with tropomyosin in the C (Ca^2+^ induced) state position (Figure S10f, g). Interestingly, the position of tropomyosin differs from that in the A-band, where tropomyosin is in the M (myosin-bound) state (Figure 4c, S10a, b). Thus, the binding of myosin to the thin filament shifts tropomyosin from the C-state to the M-state position in the A-band, while tropomyosin remains in the C-state in the I-band, suggesting that tropomyosin position can vary on a local scale within the same sarcomere and even thin filament.

### The organisation of the Z-disc

The irregular pattern of thin filaments resumes to an ordered state when the filaments approach the Z-disc (Figure 4g). In contrast to the exact square patterns observed in Z-discs from midshipman fish sonic muscle^41^ and rat soleus and cardiac muscle^42^, the Z-discs in our reconstructions are less well ordered and form squared to more rhomboid patterns (Figure 4g, h, S11). This is likely due to the fact that we used single myofibrils for our studies instead of complete muscles, where Z-discs are laterally anchored and stabilized. Antiparallel thin filaments in the Z-disc with opposing polarities are connected by 33 nm long cross-links which we attribute to α-actinin^10,43^ (Figure 5a-c, S11b).

**Figure 5.**
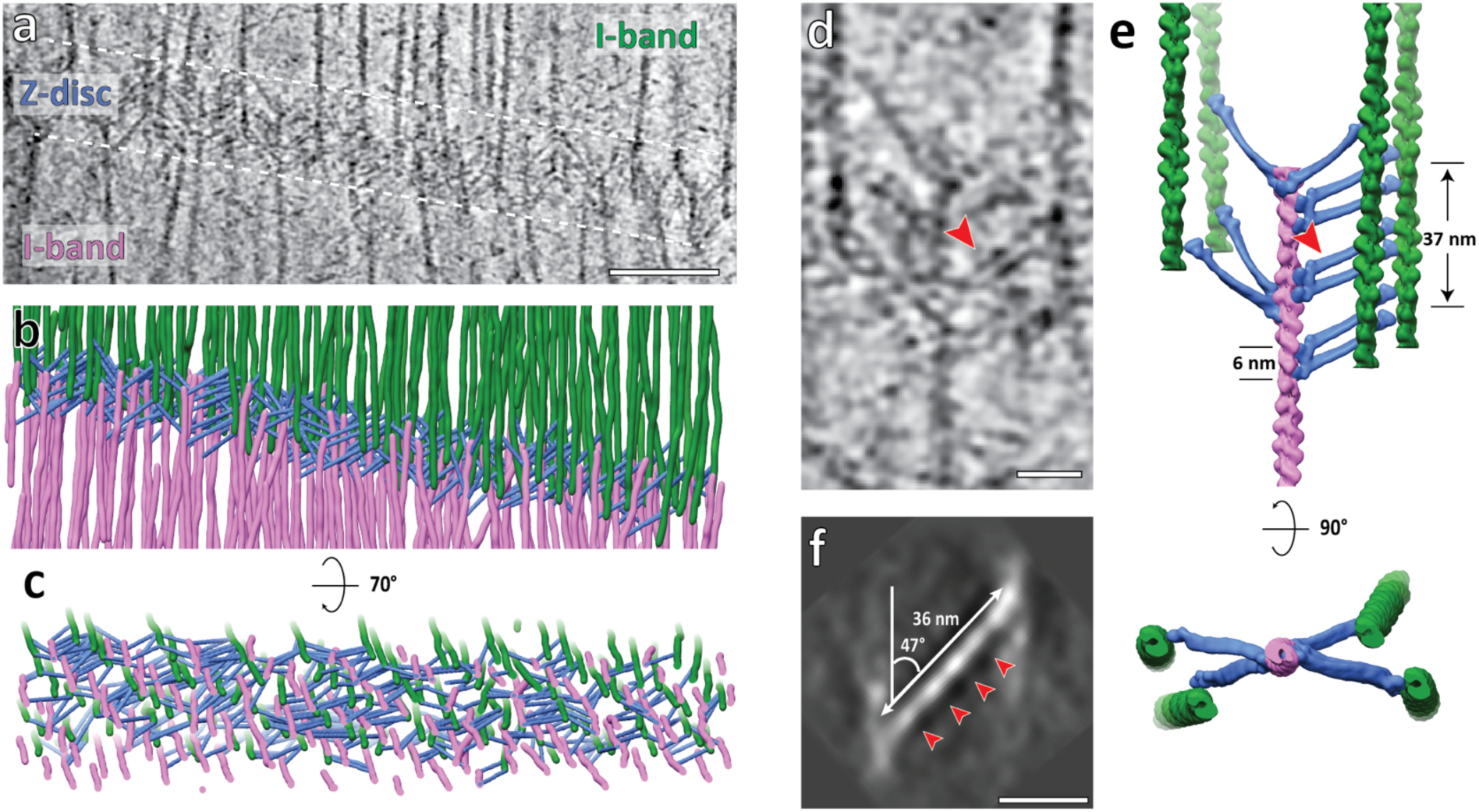
α-Actinin organisation between antiparallel thin filaments in the Z-disc. **a**, Slice through a tomogram depicting the Z-disc and two I-bands. Image was taken from a SIRT-like filtered tomogram further filtered by a low-pass filter. Scale bar, 100 nm. **b**, Cryo-ET based 3D model of the Z-disc showing antiparallel thin filaments from two adjacent sarcomeres (green and magenta) and the α-actinins (blue) cross-linking the filaments. **c**, Tilted view of the model shown in (**b**). **d**, Slice of an example unit in the Z-disc depicting one thin filament with its neighbouring antiparallel thin filaments and the α-actinins connecting them. A doublet of α-actinins is highlighted by a red arrow head. Scale bar, 20 nm. **e**, Cryo-ET based 3D model of the same region in (**d**). Actin filament model is derived from the thin filament reconstruction in Figure 4a, excluding tropomyosin. The α-actinin model is derived from the crystal structure of an α-actinin dimer (PDB: 4D1E). **f**, Projection image (7 nm thickness) of the 3D reconstruction of α-actinin obtained from averaging the sub-volumes as picked in (**b**) and (**c**), depicting four domains (marked by red arrow heads) which correspond to the four spectrin-like repeats (SRs) in the rod region. See also Figure S12. Scale bar, 20 nm.

By comparing Z-discs from different reconstructions, we found two distinct types of Z-discs. One appears to be thicker than the other (∼100 nm versus 80 nm) and is at the same time more compact (thin filaments are closer laterally) (Figure S11d, e). Interestingly, while the mean length of α-actinin is similar in both types of Z-discs (Figure S12b), the average angle between α-actinin and actin filaments in direction to the pointed end differs considerably; ∼152° in the thick form and ∼128° in the thin form (Figure S12a).

We believe that the two different forms of Z-discs represent two states of the sarcomeric unit. The thick form resembles a possible state in which the actin filaments are under low strain and the connecting α-actinins are at a more acute angle (Figure S11c). The thin form represents the Z-disc under high strain. The angle between actin and α-actinin becomes more obtuse and the Z-discs fold together like a parallel hinge or umbrella (Figure S11f). Considering a stable interaction interface between the actin-binding domain of α-actinin and actin filaments^44^ and the rigidity of the rod region of α-actinin^10^, the flexibility in the neck between the rod and ABD of an α-actinin probably serves as the central hinge of this pivot-and-rod structure^45^.

We annotated all clearly visible densities in the thinner Z-disc where single α-actinin densities were more apparent to understand the α-actinin organisation between antiparallel thin filaments (Figure 5a, Movie S2). Top and side views of the resulting three-dimensional model reveal that α-actinin forms a mesh by cross-linking the actin filaments, which is less well ordered than anticipated from the projection (Figure 5a-c) or from idealised helical reconstructions in previous work. A prominent feature is a doublet of α-actinin cross-links with 6 nm spacing, which is only possible when two α-actinins bind to longitudinally adjacent actin subunits along the actin filament (Figure 5d, e, S12c, d). We observe up to three of these α-actinin pairs connecting two antiparallel thin filaments over ∼37 nm, at ∼18.5 nm intervals. Doublets have also been observed in an *in vitro* reconstituted α-actinin-F-actin raft^46^. Our analysis shows that this arrangement is not an *in vitro* artefact but a feature of native sarcomeres.

Consistent with previous studies^41,47^, we observed several α-actinin cross-links ∼37 nm spaced (Figure S12c, e), which corresponds roughly to the half-helical pitch of actin filaments. However, we also observed α-actinins that do not follow this pattern (Figure 5d, e, Figure S12c), suggesting a less ordered organisation the Z-disc than previously assumed or inferred from atypical actin-α-actinin arrays in nemaline rods or midshipman muscle^41,48^. Though limited in Z-views of α-actinin, by performing sub-volume averaging of the α-actinin positions that we determined we could show that it connects opposite sides of antiparallel actin filaments, forming a slight “S” shape (Figure S12f). The actin binding domains and the central rod region of an α-actinin from the crystal structure^10^ fit well into the density (Figure S12f) and the spectrin-like repeats can be distinguished in a projection of the reconstruction (Figure 5f). This “S” shape is similar to the conformation of α-actinin in the crystal structure^10^, but differs from the conventional “basket-weave” model which is derived from observations of transverse sections of Z-discs^49^. Future studies at higher resolution will enable to elucidate the molecular basis of this interaction and shed light on the organisation of other proteins in Z-disc and their roles in signalling^50,51^.

The cryo-ET study of skeletal muscle sarcomeres reveals unexpected conformational plasticity of myosin heads, actin filaments, actomyosin cross-bridges and Z-disc cross-links that will inform future modelling approaches and aid in reinterpreting the intermolecular interactions of sarcomeric sub-compartments.

## Acknowledgements

We thank M. Saur for preliminary work and J. Mahamid and J. Plitzko for advice on cryo-FIB milling. We are thankful to S. Tacke for hardware optimisation for cryo-FIB and R.S. Goody for critical reading of the manuscript. This work was supported by funds from the Max Planck Society (to S.R.), the Wellcome Trust (Collaborative Award in Sciences 201543/Z/16/Z to S.R. and M.G.), the European Research Council under the European Union’s Horizon 2020 Programme (ERC-2019-SyG, grant no. 856118 to S.R. and M.G.) and the Medical Research Council (MR/R003106/1 to M.G. and A.L.K.). M.Gr. was supported by an EMBO Long-Term Fellowship. M.G. holds the BHF Chair of Molecular Cardiology. We are grateful to O. Hofnagel and D. Prumbaum for EM support and B. Brandmeier for technical assistance.

## Author Contributions

S.R. designed and supervised the project. A.L.K. and M.G. isolated mouse psoas myofibrils. Z.W. prepared cryo-lamellae, collected cryo-ET data and performed sub-volume averaging. M.Gr. optimised cryo-ET data acquisition. Z.W. and M.Gr. annotated and analysed the A-band and I-band data. Z.W., M.Gr. and T.W. analysed Z-disc data. T.W. implemented tools for data analysis and automatic filament picking. Z.W. and M.Gr. prepared figures and videos. Z.W., M.Gr., and S.R. wrote the manuscript. All authors reviewed the results and commented on the manuscript.

## Author Information

The coordinates for the EM structures have been deposited in the Electron Microscopy Data Bank under accession numbers …… Correspondence and requests for materials should be addressed to S.R.

## Ethics declarations

Animals were sacrificed in a schedule-1 procedure by cervical dislocation following licensed procedures approved by King’s College London ethics committee and the Home Office UK.

## Methods

### Myofibril preparation and vitrification

Myofibrils were isolated from BALB/c mouse psoas muscle essentially as previously described^52,53^. Briefly, bundles of the freshly excised *psoas major* muscle were tied to plastic supports and their sarcomere length adjusted as judged by laser diffraction to 2.2 - 2.4 µm. They were allowed to equilibrate in rigor buffer (20 mM HEPES pH 7, 140 mM KCl, 2 mM MgCl2, 1 mM EGTA, 1 mM DTT, Roche complete protease inhibitor) over night at 4 °C. The following day, the central section of the bundles was dissected into ∼2 mm pieces and homogenized 3 to 4 times at 7,700 rpm rpm for 15 s, and 3 times at 10,000 rpm for 5 s (IKA TC10 basic ULTRA-TURRAX® homogenizer with S10N-5G dispersing element, IKA England) with intermittent washes in rigor buffer, with myofibril separation monitored by light microscopy of small samples at each step. Myofibril suspensions were stored in complete rigor buffer at 0 °C for 1-3 days until vitrification. Grids with myofibrils were vitrified using a Vitrobot Mark IV plunger (Thermo Fisher Scientific, USA); 2 µl of myofibril-containing solution was applied to glow-discharged Quantifoil R1.2/1.3 Cu 200 grids. After incubation on the grid for 60 seconds at 13 °C, excess solution was blotted for 15 seconds from the opposite side from the sample with a Teflon sheet attached to the front pad of the apparatus. The grids were then vitrified by plunge-freezing into liquid ethane and afterwards clipped into cryo-FIB-specific autogrid rings (Thermo Fisher Scientific) with marks for orientation alignment and a cut-out for subsequent milling at a shallow angle.

### Cryo focused ion beam milling

Clipped autogrids were loaded into a shuttle and transferred into an Aquilos cryo-FIB/SEM dual-beam microscope (Thermo Fisher Scientific). Overview tile sets for the grids were acquired at 256 x magnification with the scanning electron beam using MAPS software (Thermo Fisher Scientific). The grids were then sputter coated with platinum for 15 seconds to minimise charging effect with electron beam, allowing better recognition of myofibrils (Figure S1). Vitrified myofibrils were localised and labelled as lamella sites. Prior to FIB-milling, organometallic platinum was deposited onto the grids through a gas-injection-system to prevent damage to the sample at the leading edge from the gallium ion beam. For each lamella site, the coincident point between electron beam and ion beam was determine by adjusting stage Z height. The stage was tilted to allow a 6-10° incidence angle of ion beam. FIB-milling using gallium ions was afterwards performed in four steps as described in Table 1. During each step, all lamellae were milled before proceeding to the next step. All lamellae were polished within one hour to minimise water deposition onto the surface of lamellae. During polishing, the lamella was monitored with the electron beam at 5 kV, 25 pA to help estimate its actual thickness via charging propensity.

**Table 1.** Milling strategy during lamellae production.

|  | Ion beam voltage | Ion beam current | Thickness of lamella |
| --- | --- | --- | --- |
| Step 1 | 30 kV | 500 pA | 3 $\mu\text{m}$ |
| Step 2 | 30 kV | 300 pA | 1 $\mu\text{m}$ |
| Step 3 | 30 kV | 100 pA | 500 nm |
| Step 4 (polishing) | 30 kV | 50 pA | 30-150 nm |

### Electron cryo-tomography and tomogram reconstruction

Autogrids were rotated by 90° after milling before being inserted into an autogrid cassette so as to align the longitudinal axis of lamellae perpendicular to the tilt axis of the transmission electron microscope (TEM) stage later. This ensures autofocus and record positions are aligned along the tilt axis on a lamella have close to the same eucentric height. Grids were then loaded into a Titan Krios (Thermo Fisher Scientific) equipped with a zero-loss energy filter and a K2 Summit direct electron detector (Gatan Inc., USA). SerialEM software^54^ was used to acquire images. Lamellae overview images were acquired at 6,300x or 8,400x magnification. High magnification tilt series were acquired at a calibrated post-energy-filter TEM magnification of 28409x (nominal magnification of 81,000x, pixel size 1.76 Å) with a dose-symmetric tilt scheme^55^ and a total dose ranging from approximately 130 to 155 e^-^/Å^2^. The stage was tilted from -60° to +60° relative to the lamella tilt angle (6 - 10°) with an increment of 3° during data acquisition. The defocus at which acquisition was performed ranged from 2 µm to 5 µm. Collected tilt movies were subsequently motion-corrected and stacked in batch using a custom python script with MotionCorr2^56^ and EMAN2^57^. Subsequent tomogram reconstruction including tilt series alignment, CTF estimation and correction, and weighted back projection were performed using the IMOD software package^58^. Platinum particles deposited onto the surface of lamellae were used as fiducial markers for alignment of tilt series where possible whereas in their absence patch tracking was used to align the tilt series.

### Sub-volume averaging of the thick filament

Tomograms were initially 4x binned and lowpass-filtered to 60 Å using EMAN2 for visualisation. To pick filaments accurately without the interference from signals from cross-bridges, an equatorial mask in the Fourier transform of XY slices of tomograms was applied. CrYOLO^59^ was afterwards employed to detect the shape of thick filaments in the XZ planes of the filtered tomogram. The detected points were then traced by a python script and generate the coordinates for thick filaments. In total, 13,700 segments of thick filaments (sub-volumes) were picked from 8 tomograms with an inter-segment distance of 105 Å. These segments were extracted from unbinned and 2x binned tomograms with a box size of 128 pixels (450 Å) using RELION^60^. Prior information of the orientations of the thick filaments in a tomogram was used to generate a featureless cylinder-like reference using PEET^61,62^. The sub-volumes were then aligned and averaged using RELION. Three strategies were used during alignment. First, the sub-volumes were aligned without any symmetry imposed and a global refinement was enabled. This resulted in aligning the missing wedge and a map with anisotropic resolution was generated (Figure S3b). Then, a C3 symmetry was imposed during refinement according to a previous structural study of the relaxed human cardiac myosin filament^63^. Although the averaged map had an isotropic resolution, the quality of the map did not improve (Figure S3c). In the end, without any symmetry imposed, only local refinement was enabled to prevent aligning the missing wedges (Figure S3d). This average has an estimated resolution of 30.4 Å based on the “gold-standard” FSC with 0.143 criterion. The map was used as a model for the thick filament in Figure 3 and S7.

### Sub-volume averaging of the actomyosin complex and fitting of atomic model

Thin filaments were picked automatically in a similar method as described above. Instead of using crYOLO, thin filaments, which appeared as dense dots from the XZ view (Figure S2), were recognised and traced by the TrackMate plug-in^64^ in Fiji^65,66^. In total, 32,421 segments (sub-volumes) were picked from 8 tomograms with an inter-segment distance of 63 Å. These segments were extracted from unbinned and unfiltered tomograms with a box size of 200 pixels (351 Å) using RELION^60^. In order to exclude low-quality sub-volumes using 2D classification, the extracted sub-volumes were first rotated to orient thin filaments parallel the XY plane based on filament orientations calculated from particle positions. Projection images along Z-axis were then generated from the central 100 slices (175 Å) of each particle and subsequently classified using 2D classification in the ISAC software^67^ from the SPHIRE package^68^. This approach, compared to using the entire sub-volume for projection, excluded signals from other filaments at the corner of the sub-volume and thus enabled a more reliable assessment of particle quality (Figure S4b). 21,130 selected good sub-volumes were aligned, averaged, and classified in 3D using RELION, based on a cylinder-like reference generated from averaging all sub-volumes after aligning the longitudinal axis of segments using PEET (Figure S4c). In each 3D class, translation and rotation parameters were modified to align the most prominent double-head of myosin (Figure S4d). After combining the modified good classes and removing duplicate particles, 18,090 sub-volumes were refined locally with a mask including the thin filament and one pair of myosin double-head. Final average has an estimated resolution of 10.2 Å based on the “gold-standard” FSC with 0.143 criterion. Local resolution was estimated in SPHIRE using the two half-maps and the mask used during averaging. The final map was sharpened with a B-factor of -200 and filtered to the nominal resolution (Figure S4f).

In order to resolve the light chain domains of myosin heads, sub-volumes centred on myosin the double-heads (shifted 90 pixels along x axis compared to original sub-volumes) were extracted. The positions of these new sub-volumes in tomograms were calculated by a python script using the original sub-volume coordinates and their alignment parameters. The new sub-volumes were sorted through 3D classification in RELION. 4,519 particles from the two good classes were refined locally with masking out the thin filament and myosin double-head (Figure S4e, g).

An initial atomic model of the *in situ* actomyosin complex was first built by rigid-body fitting of actin subunits and myosin motor domains from the atomic model of the *in vitro* actomyosin complex^7^ (PDB: 5JLH), tropomyosins from the atomic model of isolated cardiac thin filament^40^ (PDB: 6KN8) and myosin lever arms from the crystal structure of rigor-like squid myosin S1^23^ (PDB: 3I5G) using ‘Fit in Map’ in Chimera^69^. Essential light chains (ELCs) and the regulatory light chain (RLCs) were fitted separately into the averaged map after removing 5 amino acids (M796-Y800) at the hinge on the α-helix between ELC and RLC. The model of RLC together with part of the heavy chain lever arm (K801-L839) was rigid-body fitted a segmented map generated by “Colour Zone” in Chimera. Based on this initial atomic model and the sequences of actin, myosin and tropomyosin in mouse fast skeletal muscle, a homology model was calculated using SWISS-MODEL^70^ (Figure 2).

### Annotation of A-band and fitting of molecular model

In order to visualise the organisation of cross-bridges between thin and thick filaments, the densities of a complete A-band were annotated manually using Amira (Thermo Fischer Scientific) on a slice by slice basis. A user-defined mask based on grey values was used during manual annotation to prevent the picking of weaker densities. The 2D annotation stacked to form a 3D segmented volume depicting the arrangement of thin and thick filaments as well as the cross-bridges formed by myosin heads (Figure 3a). During sub-tomogram averaging, a structure of thin filament fully decorated with myosin heads (Figure S4c) was obtained prior to 3D classification. This was low-pass filtered to 17 Å and extended to a length of 1,755 Å (∼63 actin subunits) based on the helical symmetry detected in the structure (−166.6° turn and 27.9 Å rise) using RELION helix toolbox. This long fully-decorated thin filament model was then fitted into each segmented volume corresponding to a thin filament with bound myosin heads using “Fit in Map” in Chimera. The myosin heads that matched segmented volume were kept and coloured depending on which thick filament they originated from while the other myosin heads were removed. This generated a complete map of myosin heads bound to thin filaments in the I-band (Figure 3b, S7, Movie S1).

The models of double-head and split-head conformations were made from initially fitting our actomyosin structure into the segmented volume (Figure 3c, d, S6). For the split-head conformation, the upper head from the double-head conformation was fitted into densities for both heads. The S2 domain from segments of a crystal structure^71^ (PDB: 2FXM) was manually placed at the interface between the two regulatory light chains for its speculated position and orientation. Surface models were generated with the “molmap” command in Chimera at a resolution of 15 Å.

### Analysis of preferred binding positions of myosin on thin filament

In order to investigate the relation between myosin binding and the orientation of their bound actin subunit relative to the neighbouring thick filament, the corresponding angle of each actin position relative to the first actin subunit in the thin filament was calculated based on actin helical parameters determined from the averaged structure. The angles where there was a myosin bound were divided into three groups depending the original thick filaments and plotted as a circular histogram. Values of angles from different filaments were combined by aligning the average of one myosin group (represented as red colour in Figure S8), resulting a final histogram representing all myosin-bound actin positions (Figure S8a).

In addition, the molecular model of each thin filament was converted to a myosin-binding profile sequence consisting of R, G, B, and E representing actin bound by myosin from three different thick filaments and non-bound actin, respectively. The sequences of all 30 annotated filaments were aligned by multiple sequence alignment using the “msa” package in R with the MUSCLE algorithm utilising a customised weighting matrix (Table 2, Figure S9a). When a thin filament was treated as a single strand, adjacent actin subunits had a huge change in orientation (166.6°) and thus very different preference for myosin binding, resulting in nonoptimal multiple sequence alignment. Therefore, each thin filament was considered as two actin strands and only one strand was used for multiple sequence alignment. The other strands were aligned using the same alignment as the first strands. The occurrence of myosin binding at each actin position was summed up for myosin from each thick filament and then used to colour actin filament models in Chimera, showing averaged hotspots for myosin binding on a thin filament (Figure S9b).

**Table 2.** Weight matrix used for multiple sequence alignment with MUSCLE.

| Score | R | G | B | E |
| --- | --- | --- | --- | --- |
| R | 10 | -1 | -1 | -1 |
| G | -1 | 0 | -1 | -1 |
| B | -1 | -1 | 0 | -1 |
| E | -1 | -1 | -1 | 0 |

To further investigate whether myosin prefers to bind at shorted distances between thin and thick filaments, the distances between thin and thick filaments at the positions where there is a myosin head bound and where myosin is absent were calculated using a python script. For each actin subunit not bound by a myosin head, three measurements were taken between the actin and three different neighbouring thick filaments. For an actin subunit that is bound by a myosin head, the distance between the actin subunit and the corresponding thick filament is considered as the myosin-bound measurement while the distances between this actin subunit and the other two neighbouring thick filaments are considered as the myosin-free measurements. All measurements were plotted in a histogram (Figure S8c).

### Sub-volume averaging of the I band thin filaments

Thin filaments in the I-band were picked automatically as described in the previous sections with crYOLO. 19,035 sub-volumes from 4 tomograms showing I-band in the field of view were aligned with calculated priors (phi and theta angles) and refined in 3D with using RELION. Helical processing was used to minimise missing wedge artefacts using the helical parameters determined in the A-band thin filament structure. This resulted to a final structure of I-band thin filament without troponin with an estimated resolution of 10.3 Å using the 0.143 criterion. The map was sharpened with a B-factor of - 600 and filtered to nominal resolution. Thin filaments including troponin was picked manually in IMOD. In total, 704 sub-volumes were aligned to an initial reference generated by averaging all sub-volumes without alignment in PEET. The averaged structure has an estimated resolution of 19.8 Å after masking. A homology atomic model of the I-band thin filament was calculated using SWISS-MODEL based on the atomic model of isolated cardiac thin filament^40^ (Figure 4a, S10f).

### Analysis of I-band and Z-disc organisation

Thin filaments in a tomogram showing both I-band and Z-disc were first picked automatically as previously described. The segments in the Z-disc were further curated manually and merged into the originally picked filaments. The polarities of the filaments were determined based on the location of the ends of the filaments. 3D organisation of these filaments was generated in Fiji and shown in Chimera (Figure 4f). To analyse the organisation of α-actinin in the thin-form Z-disc, potential positions of α-actinin were sampled between ends of filaments with opposite polarities. The exact location of α-actinin was searched for by aligning sub-volumes extracted at these positions to a cylinder using a limited search range with PEET. The new positions and the corresponding orientation of sub-volumes representing the location of α-actinin were clustered based on the determined new position. Obvious false-positive and false-negative positions were then curated manually. In the thick-form Z-disc, 12 α-actinins were picked manually in IMOD. The angles between the pointed end of actin and the long axis α-actinin and the lengths of α-actinins in both types of Z-discs were calculated and plotted as histograms using python scripts. The 3D organisation of the thin-form Z-disc was generated in Fiji and shown in Chimera (Figure 5b, c). A local 3D model of one thin filament with its neighbouring thin filaments and α-actinins was generated by fitting models of actin filament and plotting back a map from the crystal structure of α-actinin^10^ (PDB: 4D1E) according to their 3-dimensional positions and orientations using tools developed as part of the TEMPy software package^72^ (Figure 5e).

Sub-volumes at the position of α-actinin were extracted from the 4x binned tomogram with a box size of 80 pixels (560 Å). An initial reference was obtained by averaging all 384 sub-volumes without refinement using PEET based on their orientation calculated from the coordinates of the α-actinin and the connected thin filaments. These sub-volumes were aligned to this reference during 3D refinement using RELION to generate a final average. The central rod domain from the crystal structure of α-actinin (PDB: 4D1E) was fitted into the averaged density manually. The actin binding domain, together with actin filaments was fitted into the density based on the atomic model of actin filament decorated by the first calponin homology (CH1) domain of filamin A (PDB: 6D8C), which shares a similar structure to the CH1 domain of α-actinin (Figure 5f, S12f).

