## Supplementary Information for "Molecular plasticity of the native mouse skeletal sarcomere revealed by cryo-ET"

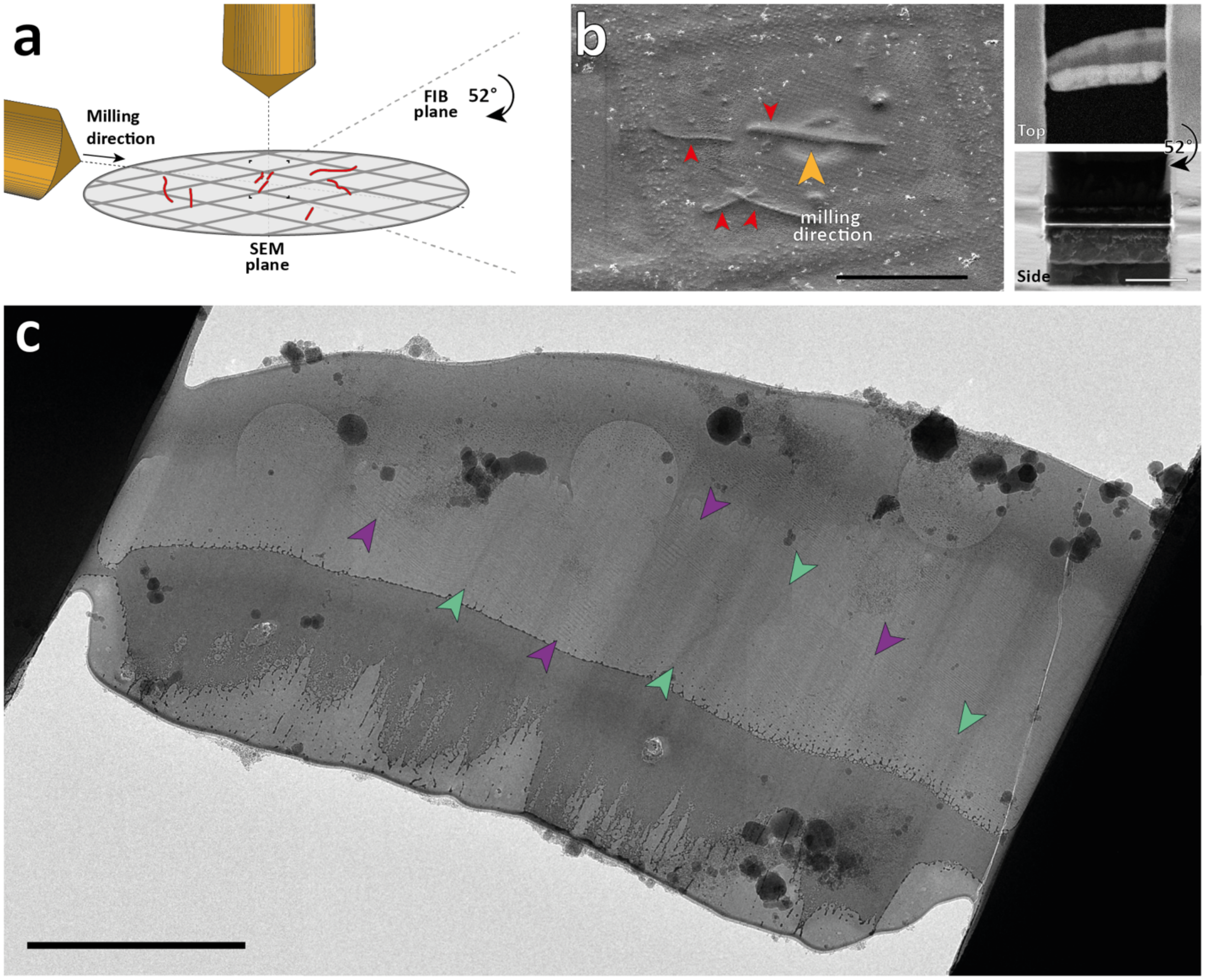


**Figure S1. Cryo-FIB-SEM of isolated mouse myofibrils. a**, Cartoon showing the arrangement of the SEM and FIB ion source with respect to the grid. Myofibrils are depicted in red. **b**, Left: SEM image taken 45° to the plane of the grid, with red arrow heads pointing toward the myofibrils. The orange arrow head indicates the milling direction. Scale bar, 50 µm. Right: Top and side views of the lamella taken after milling in the FIB-SEM. The milling angle used was 8°. Scale bar, 5 µm. **c**, Projection image of a lamella after FIB-milling recorded at a TEM. Purple arrow heads indicate M-bands of sarcomeres, while green arrow heads indicate Z-discs. Scale bar, 1 µm.

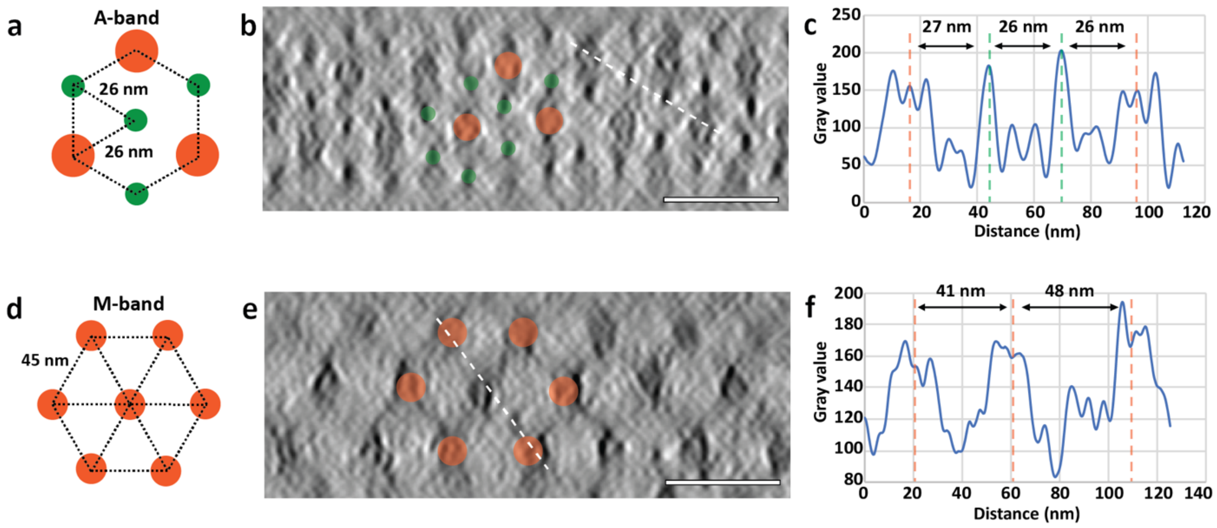


**Figure S2. Cross-section tomographic view of the mouse psoas sarcomere, showing the organisation of the thick and thin filaments**. **a, d**, Schematic diagrams of the triangular arrangement of thick filaments in the A-band (**a**) and the hexagonal arrangement of thin filaments in the M-band (**d**). Thin and thick filaments are represented by green and orange circles, respectively. **b, e**, Slices of the cross-section view of tomograms in the A-band (**b**) and the M-band (**e**). The tomograms were previously filtered with an equator filter on XY slices in Fourier space to remove signals from cross-bridges. **c, f**, Distances between filaments were measured using the line profiles of the white dotted lines in (**b**) and (**e**). Scale bars, 50 nm.

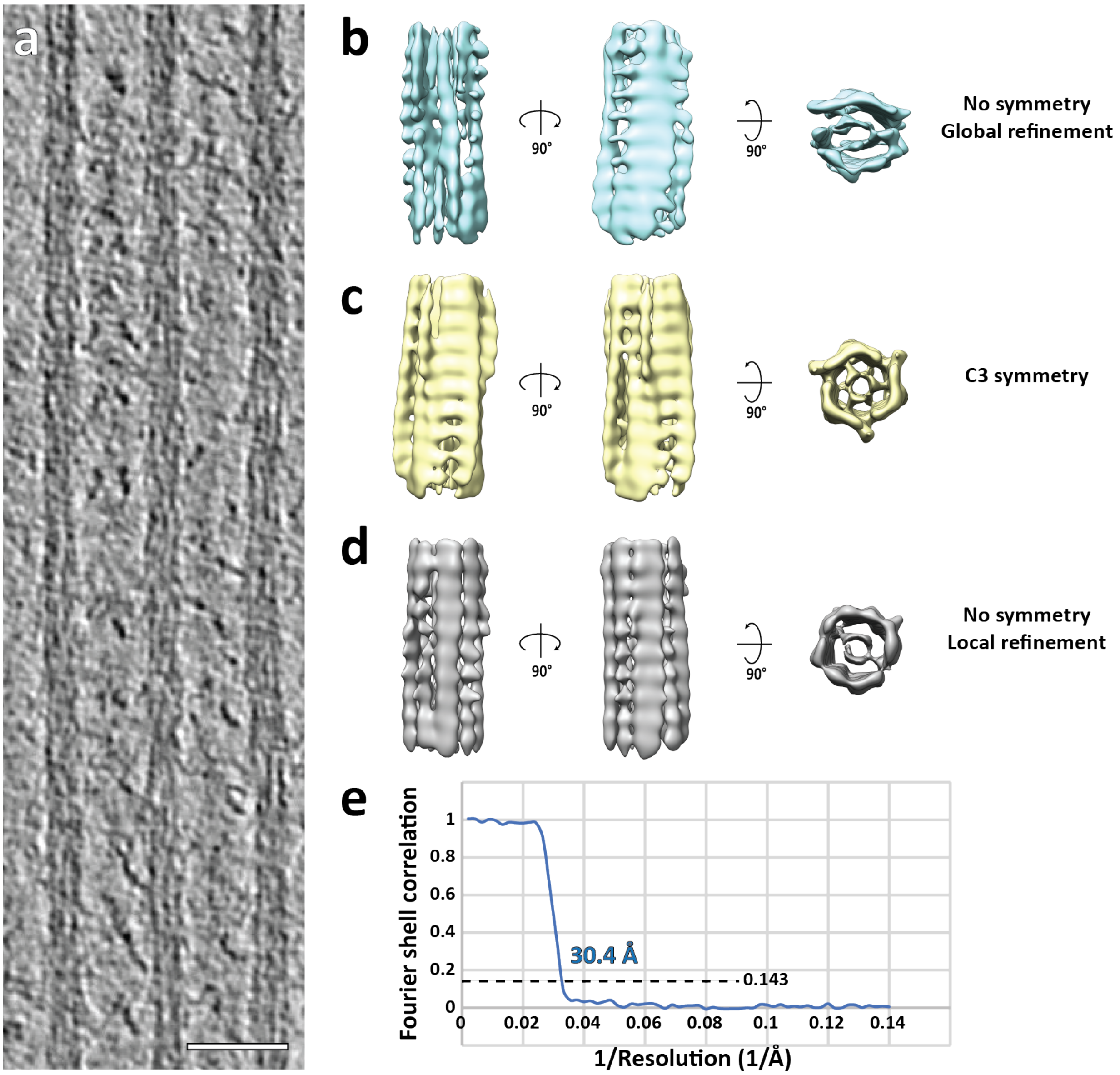


**Figure S3. Sub-volume averaging of thick filaments.** **a**, Slice through a tomogram depicting three adjacent thick filaments. Scale bar, 50 nm. **b**, Averaged structure from a global refinement. The map suffers from missing wedge artefacts, indicating alignment problems. **c**, The distribution of myosin heads when retracted to the thick filaments in the relaxed state has a C3 symmetry. We therefore performed a refinement and reconstruction applying C3 symmetry. However, this did not improve the quality of the map, indicating that the core of vertebrate thick filaments is not C3 symmetric. **d**, A local refinement with restricted possible rotation angles reduced alignment errors and thereby missing wedge artefacts. This reconstruction was used as a model for thick filament in Figure 3 and S7. **e**, The estimated resolution of the reconstruction in (**d**) is 30.4 Å using the 0.143 criterion.

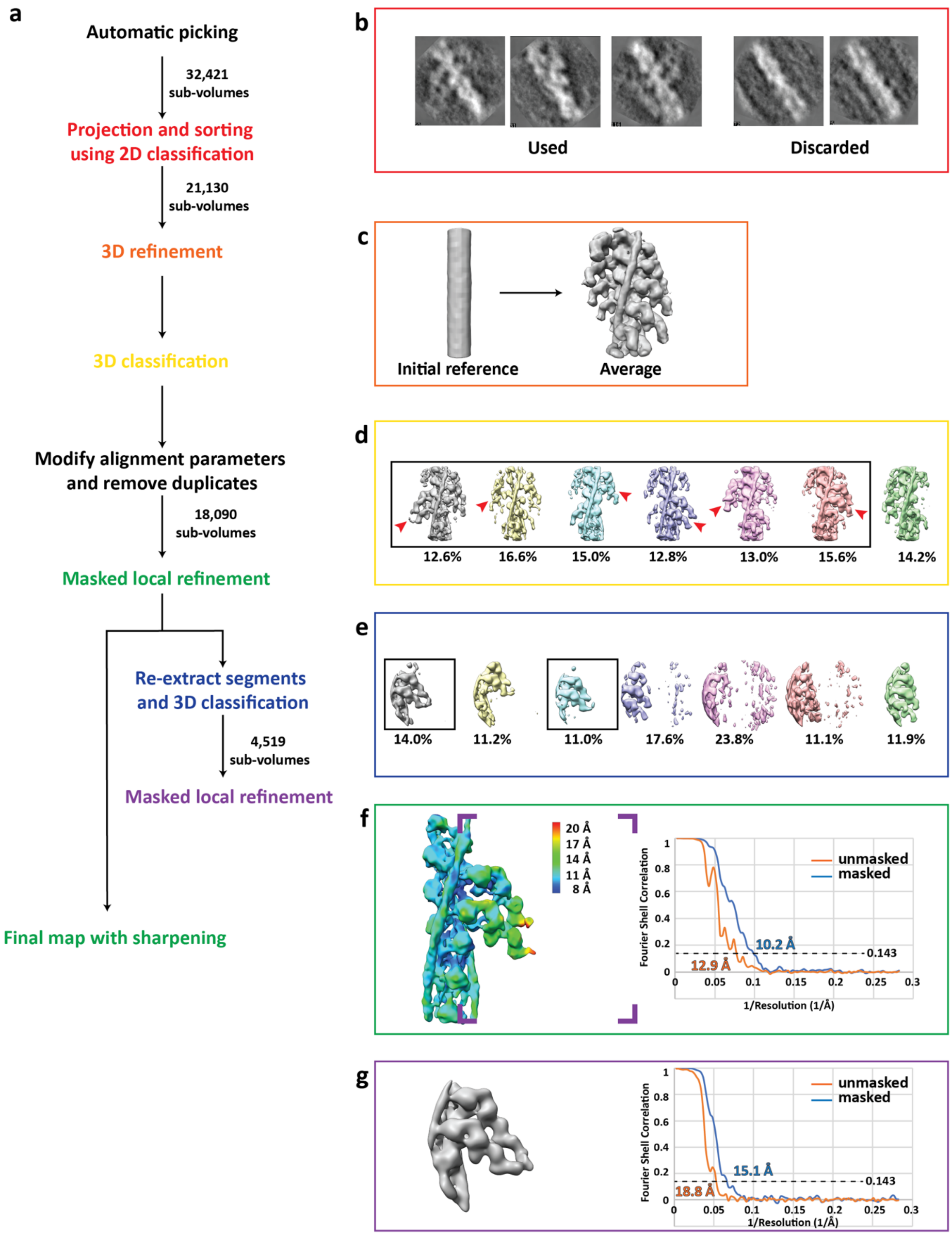


**Figure S4. Strategies of the sub-volume averaging of the A-band thin filaments. a**, Workflow of sub-volume averaging leading to the final structures of the *in-situ* actomyosin complex and double-head myosin. Details of the coloured steps are shown in (**b-g**). **b**, Examples of selected and discarded 2D class averages. After cleaning by 2D sorting, 21,130 out of 32,421 sub-volumes were selected for subsequent processing. **c**, Initial reference and the average after the first 3D refinement with only a spherical mask with a diameter of 340 Å applied. **d**, Classes of the actomyosin complex after 3D classification. Prominent myosin double head pairs in each class are indicated by red arrow heads. These pairs were later aligned to each other during merging the classes by modifying the alignment parameters. The classes in the black box were selected and used for subsequent processing. **e**, Classes of the myosin double-heads after the 3D classification applied on the sub-volumes re-centred on myosin heads. The two classes in the black boxes showing clear densities for the double-head were selected for the final averaging. **f**, Left: final reconstruction of the actomyosin complex generated from the local refinement with a mask around the thin filament and a pair of myosin heads after merging the classes in (**d**). The map is coloured based on local resolution. The purple inset shows the area of the re-centred sub-volumes for double-head analysis in (**e**) and (**g**). Right: The estimated resolution of the reconstruction is 10.2 Å using the 0.143 criterion. **g**, Left: final average of the complete myosin double-head including RLC. Right: The estimated resolution of the reconstruction is 15.1 Å using the 0.143 criterion.

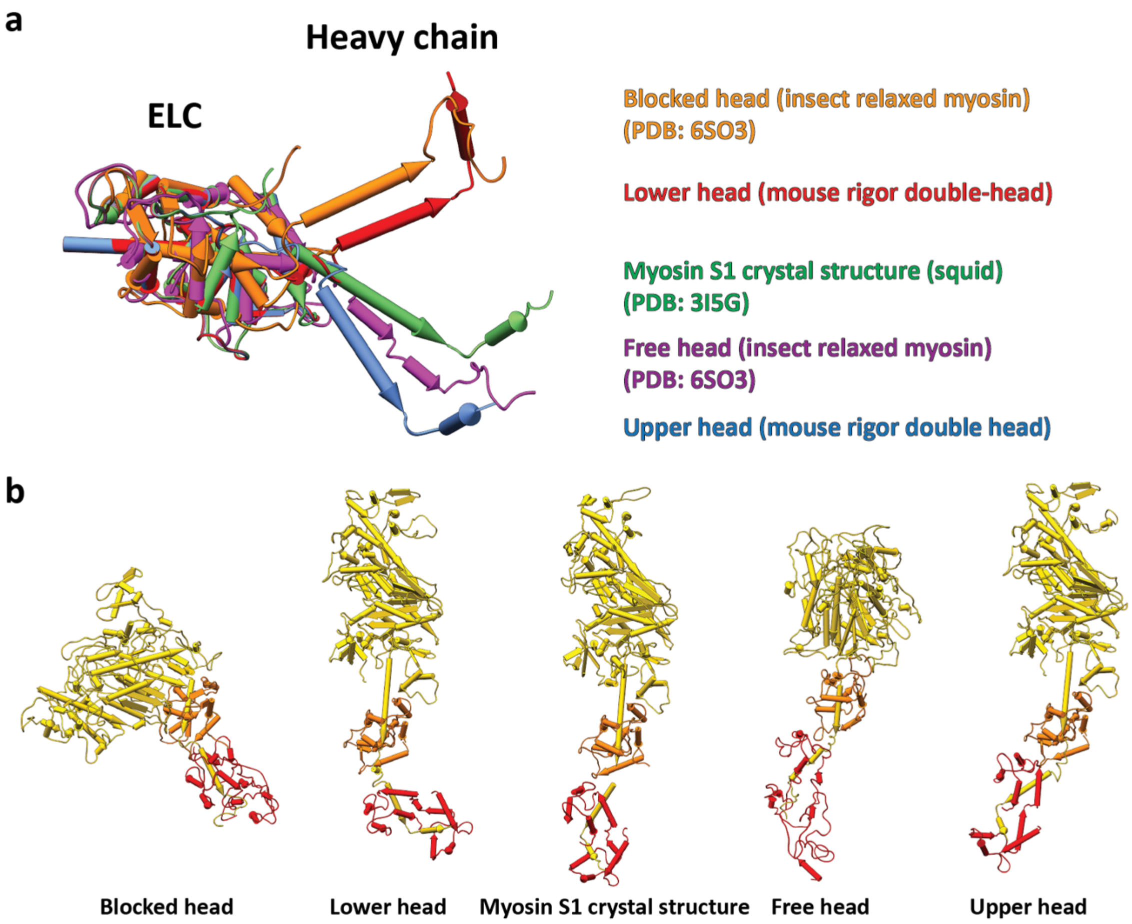


**Figure S5. Comparison of different "kink" conformations in the lever arm between ELC and RLC. a,** Alignment of ELCs, together with the ELC-binding region, from different atomic models of myosin heads. Only the RLC-binding helix of the heavy chain in each model is shown instead of the complete RLC for visualisation. Two different groups of conformations are shown. The conformation of the lower head is similar to the relaxed blocked state. The upper head exhibits a similar conformation to the relaxed free head and the crystal structure of squid myosin S1. **b**, Side-by-side comparison of the complete myosin structures in (**a**) showing both light chains.

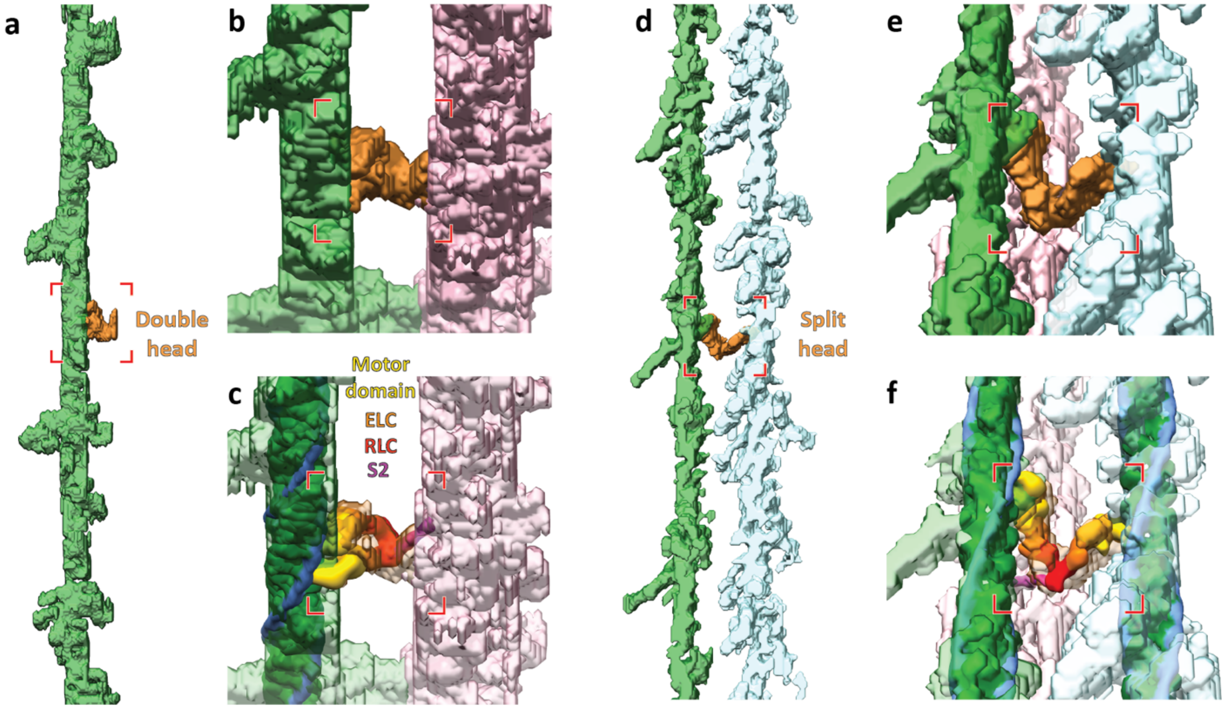


**Figure S6. Annotation and fitting of split and double heads between thin and thick filaments from within tomograms.** **a**, Annotation of a thin filament including the myosin heads (green), with the annotated density for a myosin double-head highlighted in orange. **b**, Oblique and zoomed view of the double-head density and its associated thick filament segmentation (pink). **c**, Same view as in (**b**) including the fitted thin filament and myosin double-head from sub-volume averaging. **d**, Annotation of two thin filaments including the myosin heads (green and light blue), highlighting annotated density corresponding split-head in orange. **e**, Oblique and zoomed view of the split-head density and its associated thick filament segmentation. **f**, Same view as in (**e**) including the fitted thin filament and myosin split-head. The split-head is composed of two heads in the upper head conformation in order to be fitted into the density.

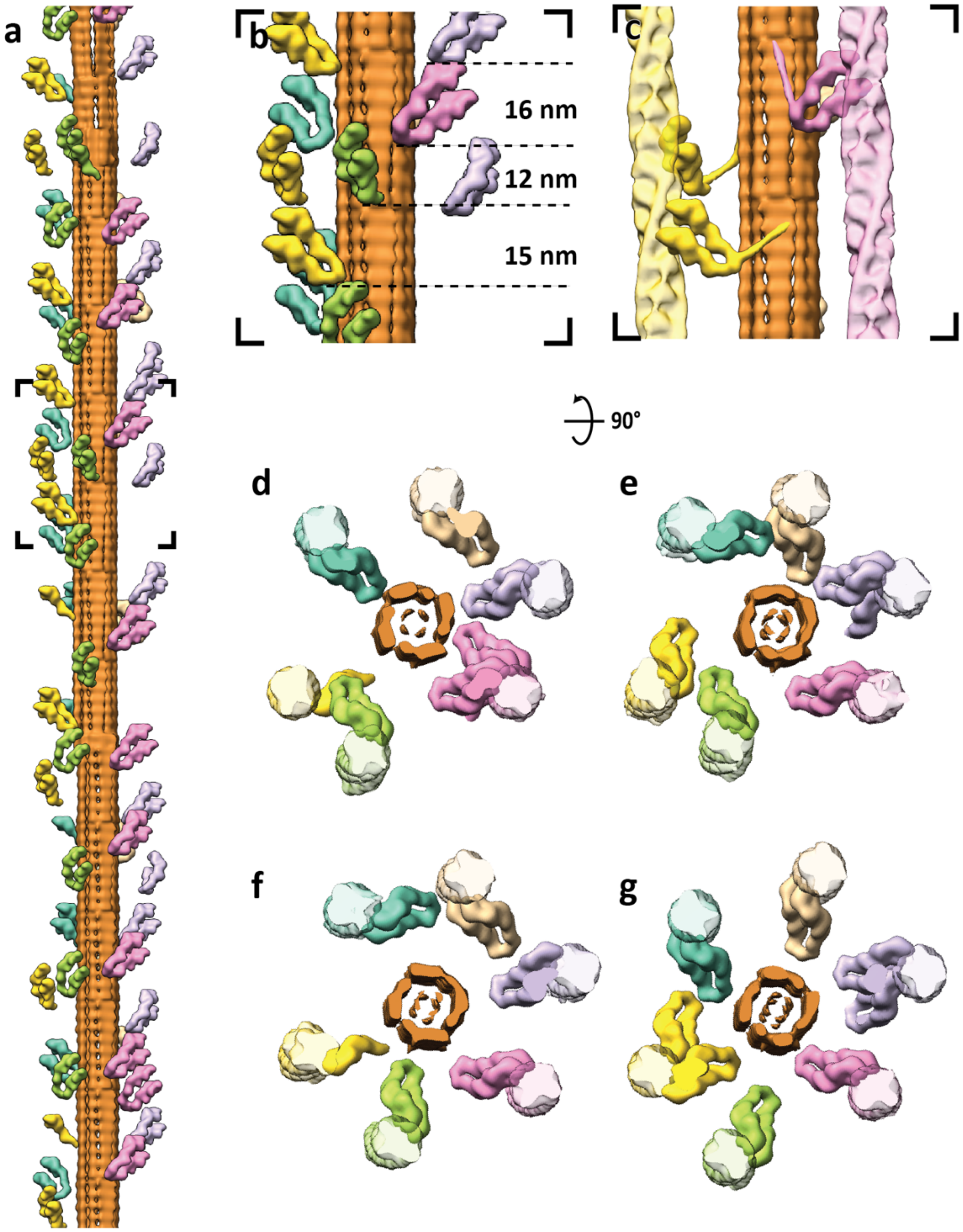


**Figure S7. Arrangement of myosin heads around a thick filament. a,** 3D model of a thick filament with surrounding myosin heads that are bound to thin filaments obtained from fitting of reconstructions from sub-volume averaging into the annotated tomogram. The six different colours represent myosin heads bound to the six adjacent thin filaments. **b,** Close-up view of the black inset in (**a**), showing varying spacings between the tips of adjacent myosin heads instead of the 14.3 nm spacing according to the helical symmetry of thick filaments. **c**, Different conformations of S2 fragments. Depending on the position of the S2 fragments in the tomogram, they are difficult to trace. Therefore, only S2 fragments that could be clearly annotated are shown. **d-g,** Top view of four segments of the thick filament in (**a**) together with the thin filaments it binds to highlighting the heterogeneity of myosin head organisation. Each image depicts a segment with a depth of ~40 nm.

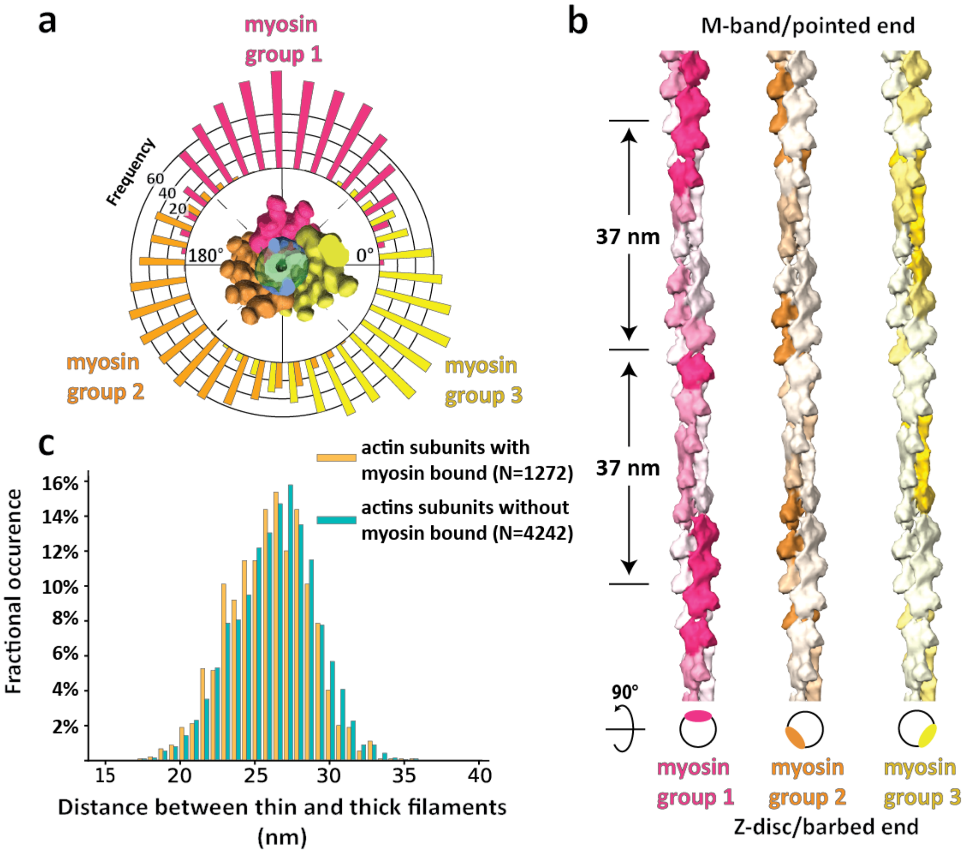


**Figure S8. Myosin binding preference dependent on actin orientation and distance between thin and thick filaments.** **a**, Angular distribution of all myosin heads shown by a circular histogram of the orientation of the actin subunits that are bound by myosin heads. The colours indicate three different thick filaments where the myosin heads originate. **b**, Hot spots of myosin binding from three different thick filaments shown on actin filaments. Actin filaments are coloured with the footprint of the myosin heads according to results of the multiple sequence alignment of the myosin binding profile sequences determined from the annotated volume and the fitted model in Figure 3b. See also Figure S9. The preferred side on a thin filament for binding of a certain myosin group is shown in the schematic diagram below. **c**, Histograms of distances between thin and thick filaments at actin subunits with (orange) or without (green) a myosin head bound. Mean distances are 25.9 nm with an SD of 2.7 nm and 26.3 nm with a SD of 2.7 nm, respectively. Skewness measures showed no skewness in either population, meaning both distributions can be considered Gaussian. The thin filament where an actin subunit is bound by myosin is on average 0.4 nm closer to the thick filaments (95% confidence interval 0.2 nm - 0.6 nm) compare to the thin filament where an actin subunit is myosin-free. This small difference implies only a minor preference for myosin binding when there is a shorted distance between the thin and thick filaments.

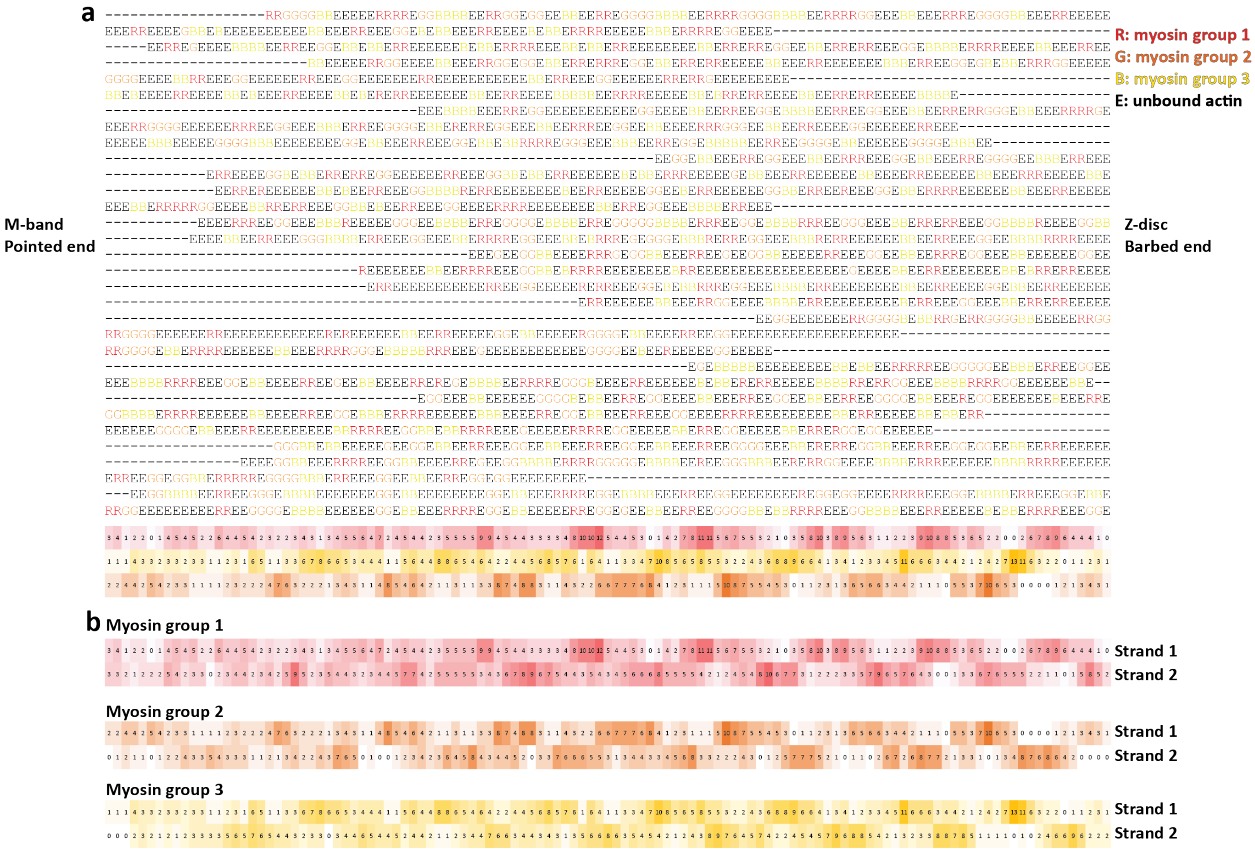


**Figure S9. Overall myosin binding profile generated from multiple sequence alignment of myosin binding profiles of 30 thin filaments.** **a**, The fitted myosin head model in Figure 3b was converted to myosin binding profile sequences on thin filaments, with "R", "G", "B" representing actin subunits bound by three groups of myosin heads originating from three neighbouring thick filaments and "E" representing myosin-free actin subunits. These sequences were aligned using multiple sequence alignment with a customised weight matrix. Only the sequence of one actin strand for each thin filament was used for alignment. The sum of occurrence of each myosin group at each actin subunit position is shown at the bottom and coloured accordingly. Darker colour indicates higher occurrence. **b**, The coloured overall myosin binding profile on both actin strands for each myosin group. Sequences of strand 2 were combined using the alignment of strand 1 shown in (**a**). The overall myosin binding profiles depict hot spots for myosin binding on a thin filament.

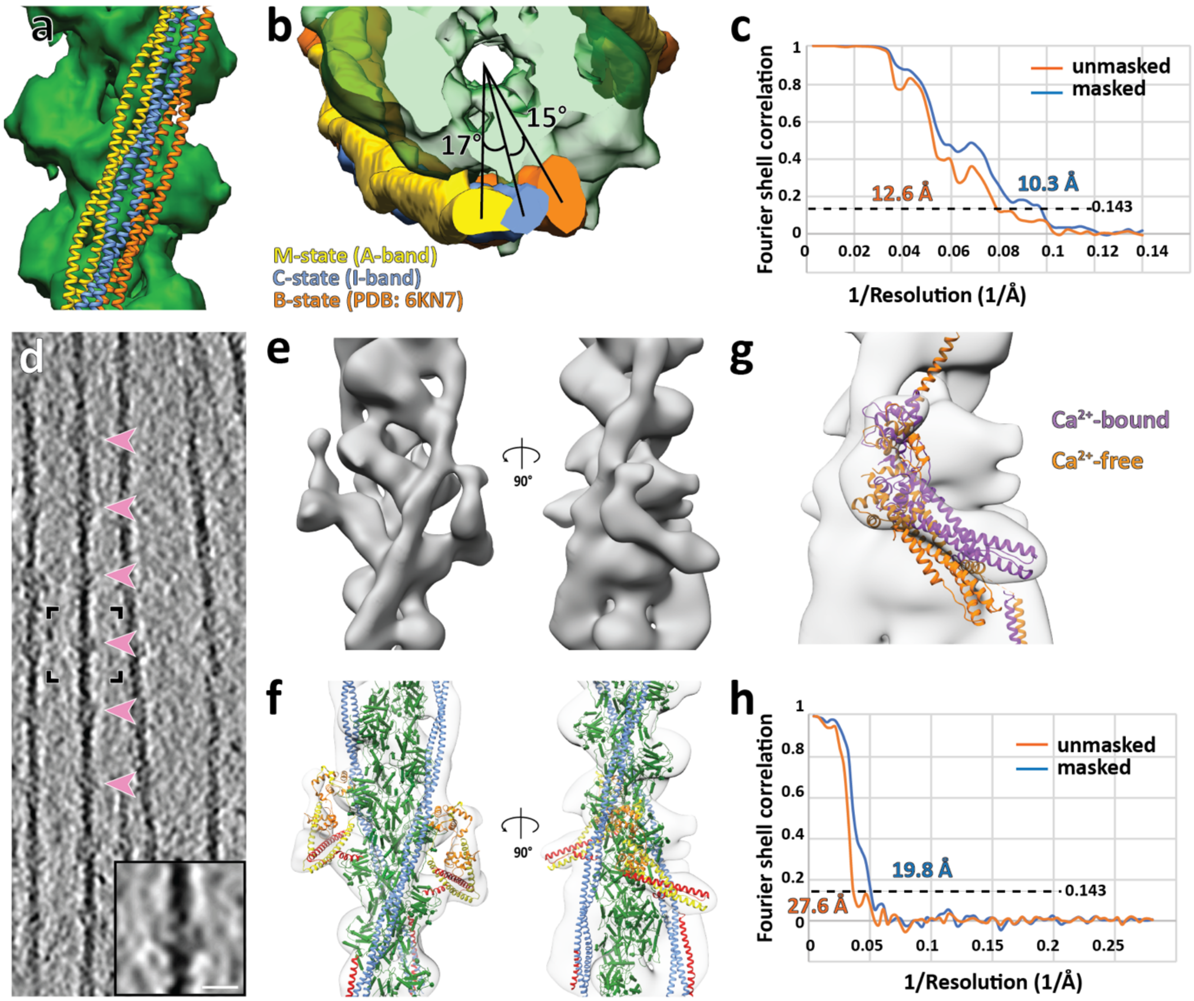


**Figure S10. Thin filaments in the I-band. a, b,** Comparison of tropomyosin positions in different regions of the sarcomere. Top and side views of a thin filament with tropomyosin in the A-band (yellow) and I-band (blue). Tropomyosin is in the M state in the A-band and resides in the C-state in the I-band. The two states differ by an azimuthal rotation of ~17°. The B-state tropomyosin from a model of $Ca^{2+}$-free thin filament (PDB: 6KN7) (orange) is also shown for comparison. **c,** The estimated resolution of the reconstruction is 10.3 Å using the 0.143 criterion. **d,** Slice of a tomogram depicting thin filaments in the I-band. Bulges along the filaments appear with a periodicity of ~37 nm (highlighted by pink arrow heads), corresponding to troponin complexes. An enlarged version of a pair of troponin complexes in the black inset is shown at the bottom-right corner, depicting clear extra densities. Scale bar, 10 nm. **e,** 3D reconstruction of a thin filament decorated by a pair of troponin complexes obtained from sub-volume averaging. **f,** A homology model based on the structure of the $Ca^{2+}$-bound cardiac muscle thin filament (PDB: 6KN8) was fitted into the map. Actin, tropomyosin, troponin I, troponin T and troponin C are shown in green, blue, yellow, red and orange, respectively. **g,** The structure of troponin in the $Ca^{2+}$-bound state (PDB: 6KN8) fits much better into the map than the structure of troponin in the $Ca^{2+}$-free state (PDB: 6KN7). Together with the C-state position of tropomyosin (**a,b**), this indicates that the I-band thin filament is in the $Ca^{2+}$-bound state. **h,** The estimated resolution of the reconstruction is 19.8 Å using the 0.143 criterion.

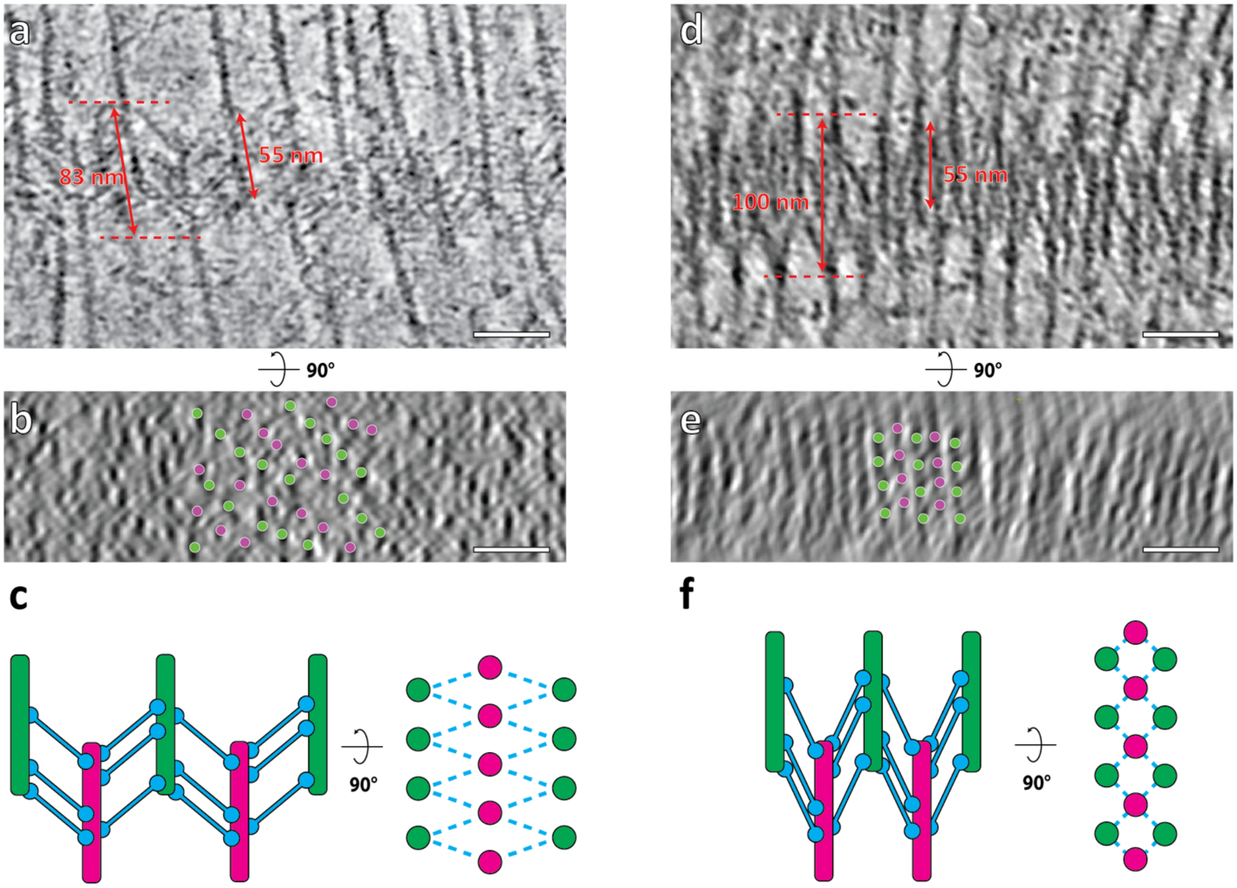


**Figure S11. Two types of Z-discs from fast mouse psoas myofibrils. a-c,** Top slice view (**a**), equator-filtered cross-section view (**b**) of a tomogram, and a schematic diagram showing the pattern (**c**) of a Z-disc in the thin form. The thickness of the Z-disc was measured between the sites where $\alpha$-actinins start to bind to the antiparallel thin filaments. The length of a single thin filament involved in the Z-disc was also measured. The organization of the filaments in this Z-disc is rhomboidal, likely resulting from a tilted orientation of the Z-disc on the grid and a slight stretching of the myofibril. We used this Z-disc for our analysis since most of the $\alpha$-actinin densities could be unambiguously annotated. **d-f,** Top slice view (**d**), cross-section view (**e**), and a schematic diagram showing another Z-disc in the thick form with a closer to square pattern (**f**). This is close to the predicted organization of a Z-disc from skeletal muscle. However, it is very difficult to recognise individual $\alpha$-actinin densities in these tomograms hampering a non-biased annotation of $\alpha$-actinins. Scale bar, 50 nm.

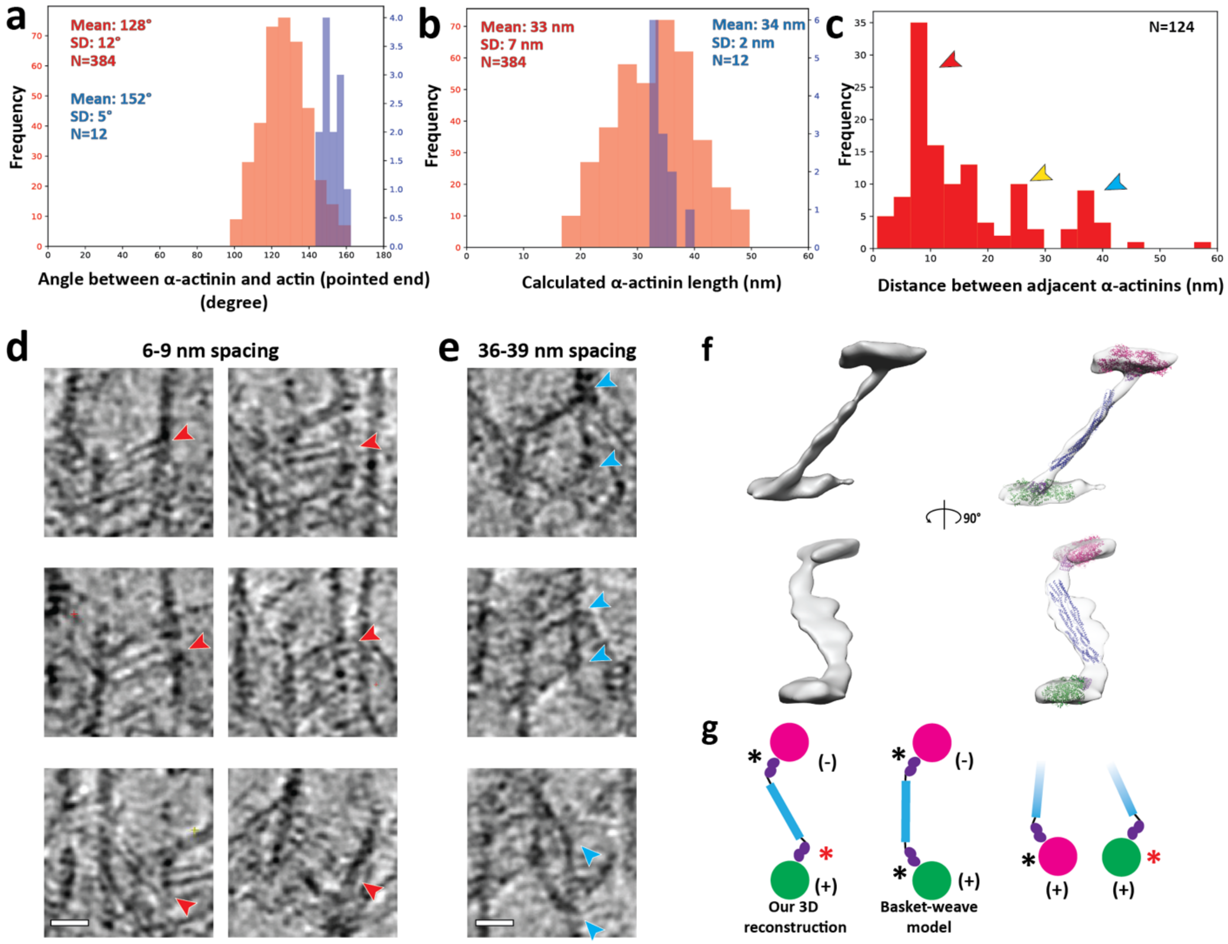


**Figure S12. α-Actinin structure and organisation in the Z-disc. a,** Distribution of the calculated angles between annotated α-actinins and actin filaments in direction to the pointed end in the thin (red) and thick (blue) form Z-discs. **b,** Distribution of the length of annotated α-actinins in the thin and thick form Z-discs. The distance along α-actinin between the centres of the connecting actin filaments was measured and the length of α-actinin was calculated by subtracting the diameter of an actin filament (6 nm). The relatively large standard deviation in the thin form Z-disc is likely caused by α-actinins binding to actin filaments at different azimuthal orientations and the error in the precise determination of the central axis of actin filaments. **c,** Distribution of the calculated spacing between adjacent α-actinins. The red, yellow and blue arrow heads highlight peaks at 6-9 nm, 24-27 nm and 36-39 nm, respectively. The 24-27 nm peak appears as a result of the two other types of spacing (36 - 2x6). **d, e,** Example images showing α-actinins with the 6-9 nm spacing (**d**) and the 36-39 nm spacing (**e**). Arrow heads highlight the positions and orientations of α-actinins. Scale bar, 20 nm. **f,** 3D reconstruction of α-actinin obtained from sub-volume averaging. Although there is a strong missing wedge effect resulting in an elongation of the reconstruction in one direction, we were able to manually fit atomic models derived from the crystal structure of α-actinin (PDB: 4D1E) and the cryo-EM structure of actin filaments bound by the first calponin homology domain of the actin binding domain (PDB: 6D8C) into the density. Actin filaments are shown in magenta and green; the actin binding domain and the rod region are depicted in purple and blue, respectively. The flexible neck regions and the C-terminal calmodulin-like domains are not shown. **g,** Left: Schematic diagram showing the difference between averaged α-actinin structure and the conventional basketweave model. Right: The two different interactions (marked by the red and black asterisks) between the end of $\alpha$-actinin and actin filaments (magenta and green) are demonstrated on the right, formed by the flexible neck region between the rod (blue) and actin binding domain (purple).

**Supplementary Video 1**

**From a tomogram to the molecular model of the A-band.** Slices of a tomogram depicting the A-band of a sarcomere are shown by moving through the Z axis. Thick (red and yellow colours) and thin filaments (green and blue colours) were annotated in 3D. Red and yellow colours represent the thick filaments. Green and blue colours represent the thin filaments, together with the myosin heads that are bound to them. The actin-tropomyosin structure fully decorated with myosin heads was then rigid-body-fitted in the annotated volume of each thin filament (grey). Myosin heads mostly outside the density were removed and only the heads that are bound to filaments are shown in the final model and coloured based on the thick filament they originate from.

**Supplementary Video 2**

**Molecular organisation of the Z-disc.** Slices of a tomogram depicting the Z-disc of a sarcomere are shown by moving through the Z axis. The traced thin filaments (green and magenta colours for filaments with opposing polarity) and annotated $\alpha$-actinins (blue) are modelled as cylindric volumes.
